# High Resolution Solvated Models Reveal Mechanisms of Allosteric Activation of mTORC1 by RHEB

**DOI:** 10.64898/2026.03.17.712089

**Authors:** Prithwish Ghosh, Arijit Maity, Kutti R. Vinothkumar, Ravindra Venkatramani

## Abstract

The mechanistic target of rapamycin complex 1 (mTORC1) is a ∼1.2 MDa dimeric assembly comprising mTOR, mLST8, and RAPTOR that integrates nutrient, energy, and stress signals to regulate cell growth. While Cryo-EM structures have provided insights into allosteric activation of the complex by the small GTPase RHEB, their limited resolution has constrained a full mechanistic understanding. Here, we combine deep learning–based AlphaFold-3 models with Molecular Dynamics Flexible Fitting and simulations to generate refined, atomistic solvated models of mTORC1±ATP±RHEB. Simulations reveal a global remodelling of the complex by RHEB, which strengthens mTOR–RAPTOR interactions while weakening mTOR–mLST8 contacts. These drive the reorganization of Kinase N- and C-lobes into a catalytically competent state in which ATP binding is stabilized enthalpically with improved Mg²⁺ coordination. Our studies present structural, energetic and dynamic changes induced by RHEB binding which collectively cause allosteric preorganization of mTORC1 for catalysis prior to substrate binding.

## Introduction

The mechanistic Target of Rapamycin Complex 1 (mTORC1) is a large (∼1.2 MDa) multi-protein complex that serves as a central regulator of cell growth and metabolic processes by sensing nutrient, energy, and stress levels. At the core of the complex is mTOR, a serine/threonine kinase enzyme. Several protein-protein interaction (PPI) based effectors modulate mTORC1 activity including activators such as RHEB (Yang et al., 2017)(Tafur et al., 2020), and inhibitors including PRAS40 (Sancak et al., 2007) and DEPTOR (Wälchli et al., 2021). Hyperactivation of mTORC1 is implicated in various cancers (Grabiner et al., 2014), prompting the development of therapeutic inhibitors. Traditional strategies have focused on blocking substrate or ATP binding. First-generation inhibitors, known as Rapalogs, target the FRB (FKBP12-Rapamycin Binding) domain and compete with substrate binding. Second-generation inhibitors compete with ATP binding at the catalytic site. However, the emergence of drug-resistant mutations—such as A2034V and F2108L (resistant to rapalogs) and M2327I (resistant to ATP-competitive inhibitors)—has driven the development of third-generation inhibitors, or Rapalinks, which combine elements of both rapamycin and ATP-competitive drugs like INK-128(Ali et al., 2022). Recently, a fourth strategy has emerged: targeting the allosteric activation pathway of mTORC1 via RHEB binding (Shams et al., 2022), which motivates us to develop a deeper mechanistic understanding of the activation of mTORC1 by RHEB.

While the centrality and the functions of the complex in physiology and cancer biology have been heavily studied since the discovery of TOR in 1991, structural information on the components of the mTORC1 complex has only emerged in the last decade(Yang et al., 2013a),(Yang et al., 2017),(Aylett et al., 2016). Cryo-EM studies first revealed that the mTORC1 complex is dimeric (**Figure 1A**) and comprises two copies each of mTOR, mLST8, and RAPTOR(Yang et al., 2017). These studies have also provided the initial structural insights into how RHEB binding alters mTORC1 conformation(Yang et al., 2017). RHEB binding brings the N-HEAT domain closer to the M-HEAT domain. This conformational change presumably shifts the FAT domain’s C-lobe, allowing the Kinase Domain (KD) N-lobe to move inward, effectively closing the catalytic cleft. This in turn repositions the ATP phosphate groups to move closer to essential catalytic residues on the C-lobe, including Mg²⁺-coordinating Asn2343 and Asp2357, and catalytic residues like Asp2338 and His2340, thereby aligning them in the “correct register” for the catalytic activity of the enzyme complex.

**Figure 1:**
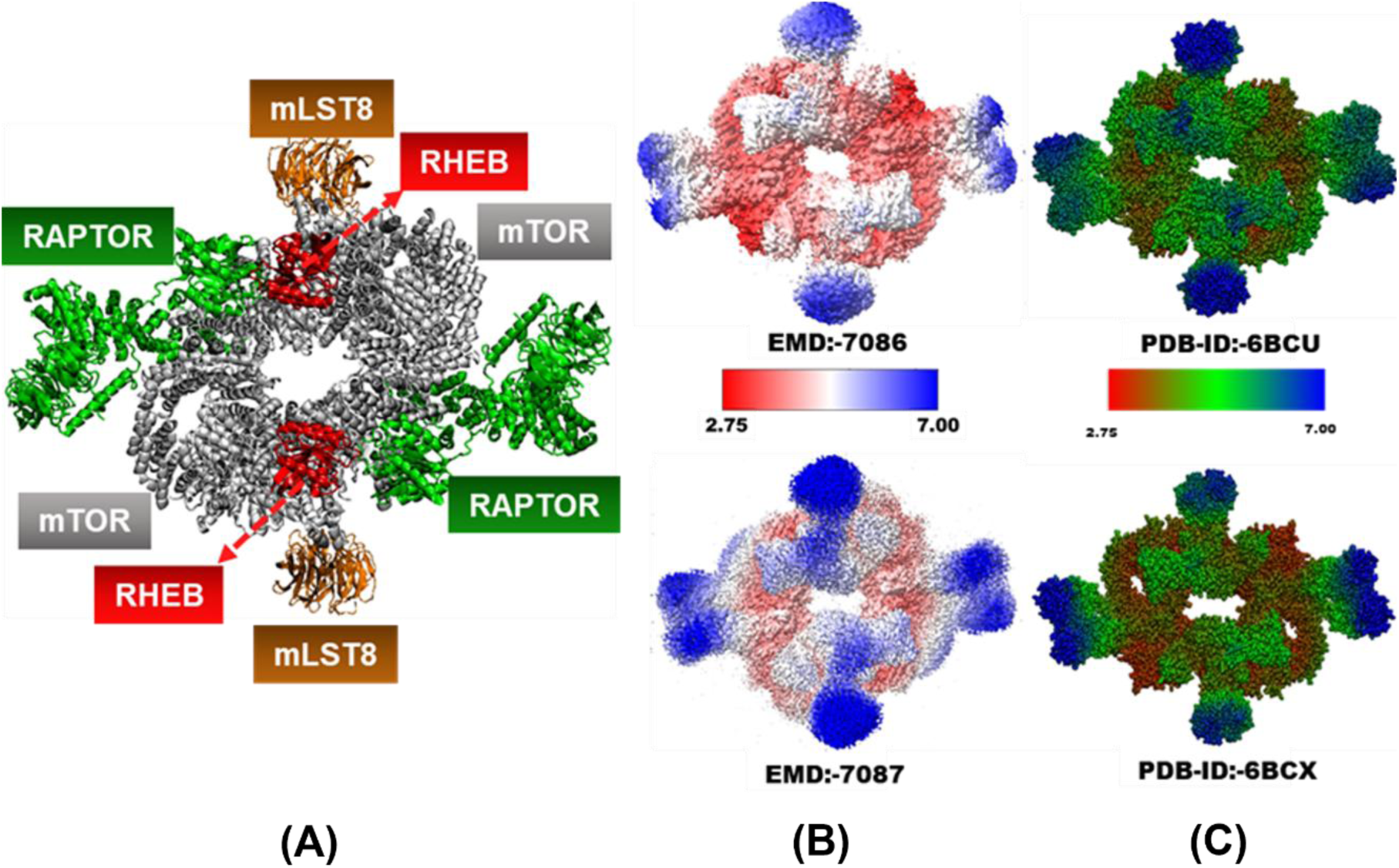
(A) Cryo-EM Structure (PBD id: 6BCU) illustrating constituent protein chains of mTORC1 with allosteric effector protein RHEB (B-C) Cryo-EM maps and Cryo-EM structures for mTORC1 complexes bound to RHEB (top: PDB id 6BCU) and RHEB-free (bottom: PDB id: 6BCX) coloured according to (B) spatial resolution and (C) atomic resolution respectively. Resolution-based colouring per atom were generated using local resolution maps using Phenix 1.21.1 (Liebschner et al., 2019) and converted to CCP4 format. Atomic resolutions were extracted from the local resolution maps by assigning each atom the local resolution at its PDB coordinate and storing the values in the B-factor column.

However, it is not straightforward to extract the details of these structural changes created by RHEB binding based on the non-uniform and low resolution Cryo-EM maps. For instance, consider the local resolution of electron density maps of RHEB-bound and RHEB-free mTORC1 in **Figure 1B** alongside their projections on the corresponding atomic models in **Figure 1C**. These maps exhibit significant resolution heterogeneity, particularly in peripheral domains where flexible loops and inter-subunit interfaces are prevalent. The spatial and atomwise resolutions also vary considerably upon RHEB binding. Moreover, the presence of unresolved (missing atoms) in the Cryo-EM model complicates efforts to study the system’s dynamics through methods such as atomistic Molecular Dynamics (MD) simulations. Such investigations can help provide a more complete view of structural, dynamical, and energetic changes induced in the complex both locally and globally, by RHEB. Finally, such models should also serve as starting points to investigate how the presence of RHEB influences substrate binding, and the phosphoryl transfer mechanism.

Traditionally, gaps in experimentally determined structures—manifesting as missing residues in coordinate files (PDB, CIF formats)—can be completed using homology-based modelling methods that rely heavily on existing structural databases. However, these approaches are inherently limited in their ability to model large segments of unresolved missing residues. This is particularly problematic for the Cryo-EM mTORC1 complexes (PDB Ids: 6BCU and 6BCX) where about 16 % of the ∼8500 residues are not resolved. On the other hand, recent deep-learning–based modelling frameworks such as AlphaFold can generate structures of individual proteins and complexes directly from sequence. The accuracy of AlphaFold2 in modelling single proteins surpassed all competing algorithms in the CASP14 experiment (Callaway, 2020)(Jumper et al., 2021), establishing it as the gold standard for single-chain structure prediction. Nonetheless, predicting multi-protein complex structures remained challenging and researchers relied on hybrid modelling strategies. For instance, a model of the SARS-CoV-2 spike protein embedded in a viral membrane was developed (Woo et al., 2020), which combined template-based modelling for loop reconstruction, ab initio monomer prediction, ab initio docking, and refinement through molecular dynamics flexible fitting (MDFF). With the introduction of AlphaFold3 (Abramson et al., 2024), accurate full-length modelling of multi-protein complexes has become feasible without depending on such elaborate workflows. However, these predictions are limited to smaller and medium sized complexes, often exhibit severe atomic clashes, and generally fail to capture large-scale conformational rearrangements associated with ligand or protein binding. The large size of the mTORC1 complex and the extensive structural rearrangements induced by RHEB binding thus cannot be directly modelled via AlphaFold and necessitate a hybrid approach.

Here we build a complete high-resolution atomistic and fully solvated model of mTORC1 with and without RHEB to interrogate local and global structural, energetic and dynamical changes induced by the effector. We used the AlphaFold server to generate initial models of the complex and refined them by fitting into Cryo-EM density maps derived from the Pavletich group (Yang et al., 2017) using Molecular Dynamics Flexible Fitting (MDFF) (Trabuco et al., 2009). Subsequently, we performed a series of energy minimization and dynamical temperature/pressure equilibration steps to resolve steric clashes and obtained starting models with high structural similarity with the Cryo-EM structure. Finally, multiple short production MD simulations were carried out on each of the refined models in the presence and absence of RHEB and ATP-Mg^2+^ to primarily demonstrate the suitability of the models towards simulations and analyse local conformational dynamics and gain mechanistic insights into the allosteric regulation of mTORC1 by RHEB. Our refined models cross-validate large scale structural changes in the N-heat and M-heat domains and reveal new insights into the changes in PPIs, ATP binding enthalpy, and complex dynamics induced by RHEB.

## Methods

### Protein Modelling & Complex Assembly

We used the AlphaFold server (https://alphafoldserver.com) (Abramson et al., 2024) to model monomeric units of the mTORC1 complex in both RHEB-bound and RHEB-free forms. The complete dimeric complex in the RHEB-bound and -free states comprise 8788 (2 × mTOR-2549, 2 × mLST8-326, 2 × RAPTOR-1335, and 2 × RHEB-184) and 8420 (2 × mTOR-2549, 2 × mLST8-326, and 2 × RAPTOR-1335) residues, respectively. In the reported Cryo-EM structures, about 16-17% of the residues were missing in RHEB-bound (PDB-ID:6BCU: Total-1428, 2 × mTOR-404, 2 × mLST8-9, 2 × RAPTOR-283, and 2 × RHEB-18), and RHEB-free (PDB-ID:6BCX: Total-1444, 2 × mTOR-430, 2 × mLST8-9, and 2 × RAPTOR-283) forms. The complexes also included ATP and two Mg²⁺ ions per chain of mTOR, along with GTP and one Mg²⁺ per chain of RHEB. While modelling the complexes, due to the sequence input limitation of maximum 5000 tokens (one token corresponds to a residue, ligand, or metal ion) on the AlphaFold server, we adopted a “divide-and-reassemble” strategy. Specifically, we first modelled each monomeric half of each complex in RHEB bound and RHEB unbound form comprising of one mTOR, mLST8, and Raptor unit. Subsequently, the full dimeric mTORC1 complex was assembled by successively aligning two copies of the monomeric AlphaFold models with each monomeric half of the cryo-EM dimer structures (PDB IDs: 6BCU for RHEB-bound, and 6BCX for RHEB-unbound) by minimizing the root mean square deviation (RMSD) of backbone *C_a_* atoms in Visual Molecular Dynamics (VMD) software (Humphrey et al., 1996).

### Molecular Dynamics Flexible Fitting to Cryo-EM data

Following the generation of the initial AlphaFold dimeric models, we removed the ligands and cofactors (ATP and two Mg^2+^within mTOR Kinase Domain, GTP and one Mg^2+^ within RHEB) and performed Molecular Dynamics Flexible Fitting (MDFF) of the modelled structures (Trabuco et al., 2008) using NAMD (Phillips et al., 2020) to the Cryo-EM derived density maps. Briefly, the MDFF protocol applies a biasing potential derived from the cryo-EM density map (**Equation Error! Reference source not found.**), driving heavy atoms of the model toward lower-density regions. The Cryo-EM density map for MDFF was obtained from the PDB structures of RHEB bound (PDB ID:6BCU) and RHEB unbound (PDB ID:6BCX) mTORC1. In VMD we use these structures to first generate simulated maps of resolution of 5Å which gets converted into an MDFF potential (𝐔_𝐄𝐌_(𝑹)) as given in **Equation** Error! Reference source not found.. Secondary structure, cis-peptide bonds and chirality restraints are also defined during the system preparation step to avoid unrealistic structural artefacts during MDFF simulations. Following this step, a rigid body docking of the AlphaFold derived dimeric models onto the maps was performed which form the starting structures for the MDFF simulations.

Multiple MDFF simulations of 50 ps each for RHEB-bound (3 runs) and RHEB-unbound (2 runs) were carried out with a time step of 0.5 fs until the RMSD of backbone atoms relative to the cryo-EM structures reaches a plateau as shown in ESI **Figure** S**1**. The larger RHEB-bound complex required longer simulations to converge the backbone-RMSD relative to the RHEB-free complex. Each of these simulations were followed by 200 minimization steps (conjugate gradient), which removes most of the atomic clashes arising while evolving the system rapidly through a biased force field. These steps included implicit solvation and were carried out after removing the ATP, GTP and Mg^2+^ from the complex. These ligands were added later to the complex (see next subsection) so that their interactions with surrounding residues matched that in the Cryo-EM structures. The medium dielectric was set to 80 to mimic a water environment.

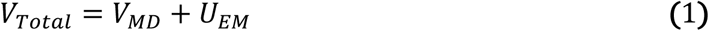

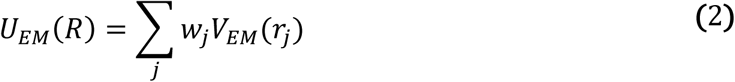

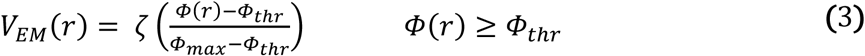

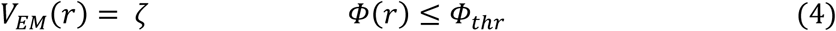

Here, the grid scaling factor 𝛇 (set to a default value=0.3 in **Equation Error! Reference source not found.**-**Error! Reference source not found.**) dictates the relative strength of the bias potential with respect to the MD potential which in this case is based on the CHARMM36 force fields(Huang & Mackerell, 2013).

### ATP Docking and Equilibration

After MDFF, ATP, GTP and Mg^2+^ were docked into the respective active sites of both the systems. Here, residues within 6 Å of ATP or GTP from the cryo-EM models were used to align the MDFF models to the corresponding regions of Cryo-EM structures, thereby preserving native ATP interactions. This procedure yielded four complete structural models (mTORC1 ± RHEB ± ATP)

We next used MD equilibration to generate fully relaxed, solvated and atomistic models of both the complexes. The equilibration protocol included sequential energy minimization, heating, and equilibration steps using NAMD. Initially each complex was placed inside rectangular water box (299.8 Å × 242.5 Å × 168.1 Å for RHEB-bound and 318.0 Å × 245.8 Å × 190.2 Å for RHEB-free complex) containing explicit TIP3 water molecules with a minimum padding length of 10 Å and the systems were neutralised using Na^+^ ions. The Charmm36 forcefield (Huang & Mackerell, 2013) was used to perform each step. All equilibration runs were carried out using an NPT ensemble and employed periodic boundary conditions. Electrostatic cut-offs were set to 16Å and all bond lengths were constrained using the SHAKE algorithm with a convergence tolerance of 1.0 × 10⁻⁸ and up to 100 iterations per timestep. Initial minimization and heating were conducted with harmonic restraints (50 kcal/mol·Å²) on heavy atoms corresponding to segments resolved in the cryo-EM structures, allowing only the modelled protein atoms (which were originally unresolved in the Cryo-EM structures) to relax. Number of conjugate gradient minimization steps is equal to 10,000 in current as well as subsequent minimization steps. Using a timestep of 1 fs, the system was gradually heated from 0 K to 300 K over the first 50 ps by reassigning velocities every 1000 steps (1 ps) in 6 K increments, followed by 50 ps of equilibration at 300 K. The minimization and heating steps were repeated with reduced harmonic restraints (25 kcal/mol·Å²). We then conducted three 150 ps equilibration stages with step size of 1fs in an NPT ensemble, gradually reducing restraints (25, 10, and 5 kcal/mol·Å²). A Langevin thermostat was employed to maintain the system at 300 K with a damping coefficient of 0.5 ps⁻¹, and pressure was regulated at 1 atm using a Langevin piston barostat with an oscillation period of 100 fs, a decay time of 50 fs, and isotropic cell coupling. Finally, an unrestrained 150ps NPT equilibration was performed keeping the same thermostat and barostat settings to yield relaxed structures, which were consistent with existing structural information as validated below. The equilibrated structures were then taken forward for production MD simulations as described in subsection 2.5 below. In summary, four systems with ∼1-1.3 million atoms were created and described in ESI **Table S1**.

### Metrics for Structural Validation

We also used several local and global metrics after each of the four modelling and simulation steps to assess how the complex structures compare with the Cryo-EM structures. We used the Local Distance Difference Test (LDDT), Global Distance Test-Total Score (GDT-TS) with cut-offs of 1-8Å, and Global Distance Test-Total Score (GDT-HA) with cut-offs of 0.5-4Å (Mariani et al., 2013) (Zemla, 2003) and (Zhang & Skolnick, 2005) and Template Modelling score (TM-score) at each stage of modelling (**Equation 5, 6**). GDT scores represent the average fraction of atoms in the model that can be superimposed onto the corresponding atoms in the reference structure within defined distance cut-offs.

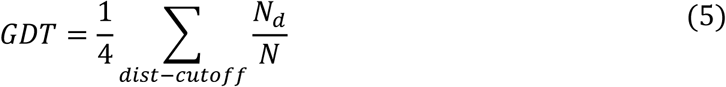

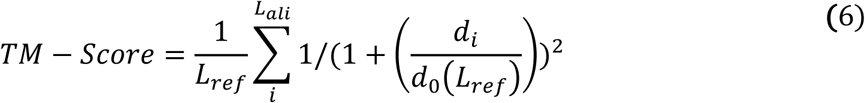

Here *N_d_* is the number of atoms within the cutoff and *N* is the total number of atoms in the reference structure. The TM-score is averaged over all aligned residue pairs between the model and reference structure

Where *d_i_* is the distance between the *i^th^* pair of aligned *C_α_* atoms, *L_ref_* is the length of the reference structure, *L_ali_* is the number of aligned residues, and *d₀*is a scale parameter that normalizes the distance for protein size. For the MD equilibrated model, we extracted 150 structures (1 structure per ps) from the final unrestrained 150 ps trajectory and extracted the mean value of these metrics with their respective standard deviations. The scores for our models are compared against distribution of best scores for TBM-Easy (TBM-Template based Modelling), TBM-Hard, FM/TBM (FM-Free Modelling) and FM from CASP13 (Guzenko et al., 2019)), CASP14 (Ozden et al., 2021)), CASP15 (Ozden et al., 2023)).

We used the TM-align software package(Zhang & Skolnick, 2005) for structural alignment, which maximizes TM-score by prioritizing structured regions over disordered regions in contrast to RMSD-based methods which treat ordered and disordered regions equally. TM-align also calculates TM-score and GDT-TS/HA for individual modelled chains with respect to Cryo-EM chains at each stage of modelling. TM-score for the full complex was calculated using MM-align (Mukherjee & Zhang, 2009) software package. LDDT calculation was performed using Open structure software package(Biasini et al., 2013). Finally, we also used the Interface Template modelling score (ITM-score) (**Equation Error! Reference source not found.**) and Interface Similarity score (IS-score) (**Equation Error! Reference source not found.**) to quantify the structural similarity of the protein-protein interface between the models and the cryo-EM structure. These were calculated using IAlign package(Gao et al., 2010).

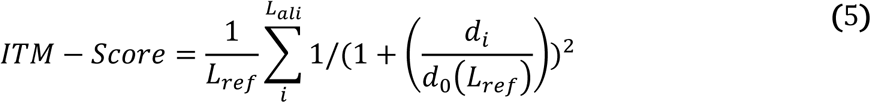

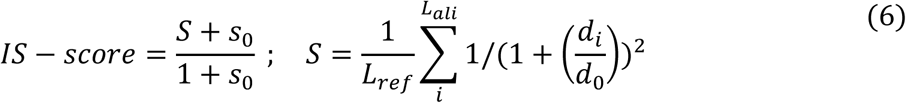

### Complex Dynamics around the MD refined Cryo-EM State

The four equilibrated models of mTORC1 ± RHEB and ± ATP were each subjected to 5×15 ns NPT production simulations to explore the local energy landscape around the refined Cryo-EM structure. Since the complexes are dimeric, statistically this amounts to 10 x 15 ns trajectories for the chains/monomers of each complex. Further for two of the models (mTORC1 ± RHEB + ATP), we extended the NPT production simulations to 5 × 25 ns to confirm and solidify our findings on the dynamics of the complexes. Here, all 5 trajectories for the complexes were initiated from the same phase space point (same coordinates and velocities) as obtained at the end of our NPT equilibration step. The production run parameters are same as the unrestrained NPT Equilibration. A comparative quantitative assessment of sampling dynamics for the complexes captured in the trajectories was then carried out using the Cumulative Variance of Coordinate Fluctuations metric (Paul et al., 2020)

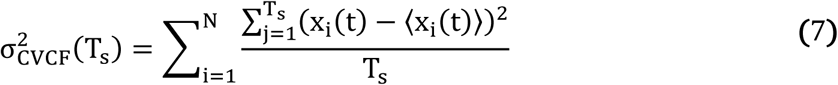

Here, *T_s_* refers to simulation time and *N* refers to number of atoms. For each set of 5×15 ns/ ×25 ns trajectories aligned at *T_s_*=0 (all trajectories all launched from the same phase space point), the variance trace is recorded as a function of 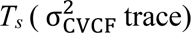. The cumulative variance was calculated for several different subsets of atoms by extracting trajectories containing said subset of atoms, which are as follows: all *C_a_* atoms of structured regions of the entire complex (5 trajectories each of mTORC1±RHEB±ATP complexes), structured *C_a_* atoms of each protein chain (10 trajectories for each chain of RHEB bound system and RHEB unbound system) and structured *C_a_* atoms of each mTOR domain (N-HEAT,M-HEAT,FAT,KD-N-Lobe,KD-C-Lobe). This constituted 10 trajectories for each chain of the four mTORC1±RHEB±ATP-2Mg^2+^systems. In each case before computing the 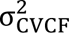, rigid body rotations and translations of the aforementioned subset of atoms were removed by aligning the atoms with respect to their positions at *T_s_*=0. The 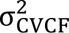 was calculated and normalized 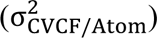 by dividing by the number of atoms *(N)*.

### ATP Binding Energetics

To quantify ATP binding to mTORC1 in the presence and absence of RHEB, we performed MM/GBSA (Molecular Mechanics/Generalized Born Surface Area)calculations (Genheden & Ryde, 2015) on the complexes with and without ATP+2Mg²⁺. We used the GBIS implementation in NAMD to perform single-point energy evaluations along the production NPT trajectories. The energy was computed over a subset of atoms comprising ATP, the two Mg²⁺ ions, and all protein residues within a range of cut-off distances (6−20 Å) from ATP. These subsets were chosen to comparatively explore distance-dependent convergence of ATP binding enthalpy to mTOR within RHEB bound and RHEB-free complexes. The total enthalpy calculated using MM/GBSA consists of the gas phase energy as well as the polar and non-polar contributions of solvation energies. We computed the change in ΔH due to RHEB binding and analysed the individual contributions to this change (ESI **Figure S7** and **Table S5**)

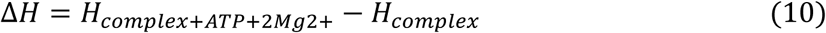

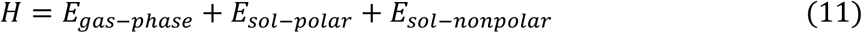

Where the distributions for ΔH were obtained over a set of 100 ±ATP-2Mg^2+^ trajectory pairs of mTOR generated by bootstrapping pairs of 10×15 ns trajectories of each mTOR unit with and without ATP. Such distributions for Δ𝐻 were calculated for both RHEB-bound and RHEB-free cases as a function of cutoff distances from the ATP as discussed above. We further generated distributions of ΔΔ𝐻 values by applying bootstrapping a set of 10,000 ± ATP-2Mg^2+^ ± RHEB trajectory pairs to estimate the average change in the ATP binding enthalpy to mTOR induced by RHEB binding to the complex. At each cutoff value we carried out the same analysis on nonbonded energies (electrostatic and van der Waals contributions) explicitly between ATP+2Mg^2+^ and the and the subset of residues lying within the cutoff.

### ATP Solvent Exposure

Percentage of Solvent Accessible Surface Area (pSASA) for the ATP was calculated as:

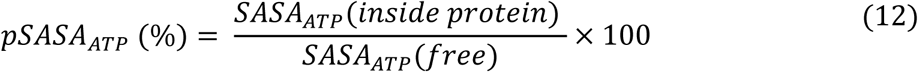

We generated both raw time traces and probability distributions of these values along the 10 × 15 ns trajectories of each mTOR chain for the RHEB-bound and RHEB-free complexes. The analysis is carried out over both the full 15 ns range of the trajectories and restricted to the last 5 ns window (where both RHEB-bound and RHEB-unbound systems exhibit stable pSASA_ATP_ values). From the distribution of pSASA_ATP_values, distinct states of ATP were identified in RHEB-bound and RHEB-free complexes. The conformational heterogeneity of ATP in each of these states was assessed by carrying out a Principal Component Analysis (PCA) of sets of structures extracted from the trajectories for which the pSASA_ATP_values lie within ± 2 % of the peaks in the distributions.

### Mg^2+^ coordination

To track the coordination sphere of both Mg^2+^ ions in MD trajectories, we looked at molecules whose oxygen or nitrogen atoms lie within a cut-off co-ordination bond distance (2.25Å,2.50Å, and 2.75Å) from the Mg^2+^ ion. The results were found to be insensitive to the value of cut-off chosen.

## Results

### Validation of Global, Local, and Interfacial Topology in mTORC1 complex models

To track the structural quality of the refined mTORC1 models during the initial AlphaFold modelling, MDFF refinement, and MD equilibration, we employed a comprehensive set of model validation metrics at the global, per-chain, local, and interface levels using the cryo-EM data as reference (Yang et al., 2017). Local structural fidelity independent of global alignment was determined using the Local Distance Difference Test (LDDT), which quantifies how well the local environment around each residue in the predicted structures is preserved relative to the cryo-EM reference (**Figure 2A**). The RHEB chains exhibited the lowest average chain-wise LDDT values (0.77 ± 0.01), yet these scores remain above the mean LDDT values reported for top-performing TBM-easy models in CASP13 (∼0.75). All other chains displayed LDDT values between 0.84 and 0.88, aligning closely with the mean TBM-easy and TBM-hard scores (∼0.87) from CASP14–15 (**Figure 2B**). These results collectively indicate that the local structural environments of our full complex models are highly consistent with those of the experimental cryo-EM structure.

**Figure 2:**
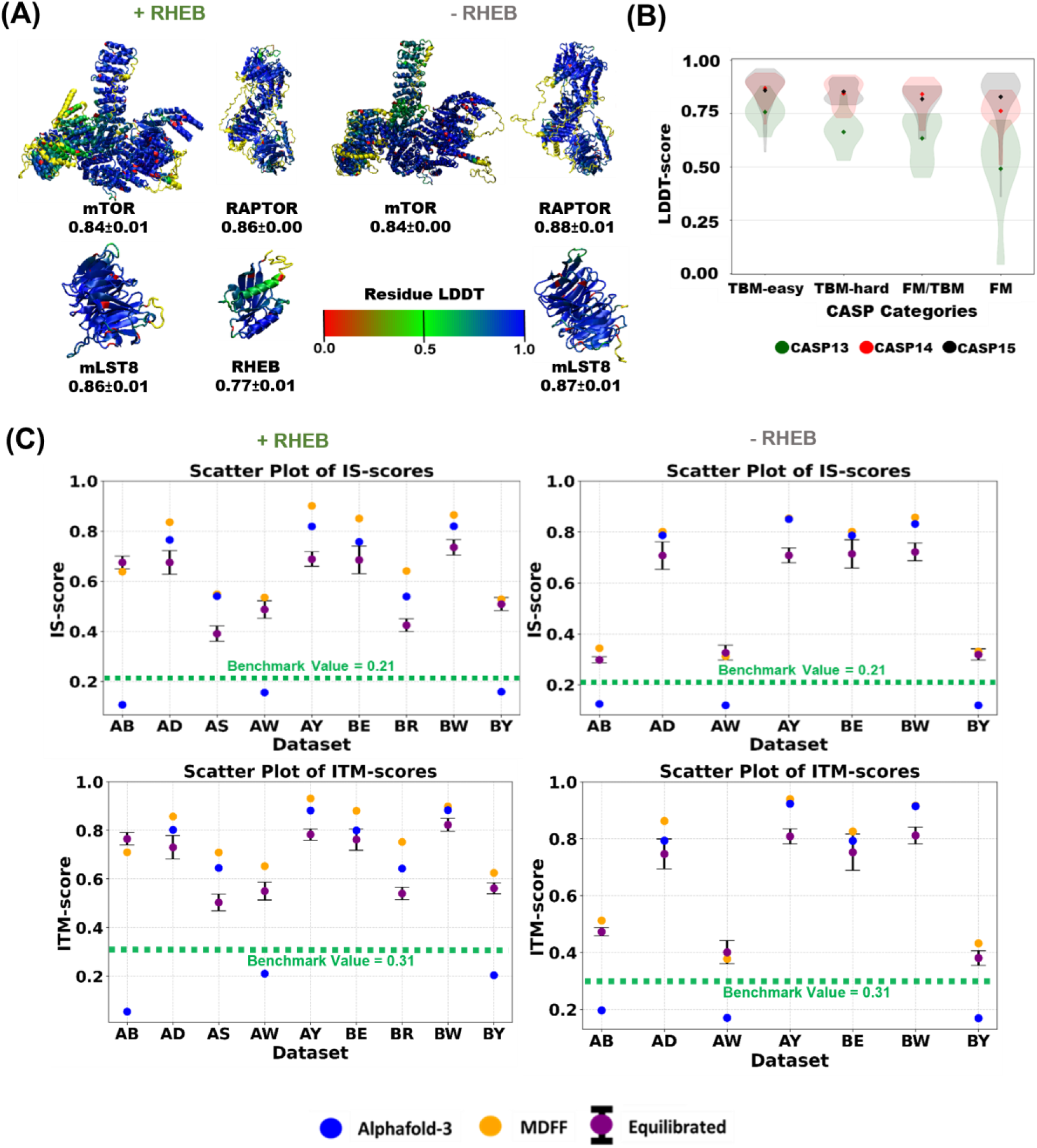
(A) RHEB-Bound and -Free models of mTORC1 chains coloured according to LDDT scores of each residue against the Cryo-EM reference structures. The regions missing in the Cryo-EM structures are coloured in yellow; (B) Violin plots representing distribution of best LDDT scores of all targets in a given category in different CASP competitions. (C) RHEB-Bound and -Free IS-scores and ITM-scores with respect to Cryo-EM reference structures along with benchmark scores for random interfaces (green dotted line).

On a global scale, the TM-score of the full complex (FC) averaged over equilibrated structures was ∼0.99 for RHEB bound and ∼0.98 for RHEB-free models, placing them in the high-accuracy range typically seen in the TBM-Easy category of CASP14 and CASP15 benchmarks. However, average TM-scores for individual proteins/chains from the equilibrated trajectories showed slightly lower values compared to FC (ESI **Figure S5** and **Table S4**). This was observed for all chains in the RHEB-Bound complex, and for mTOR (A/B) and mLST8 (D/E) chains in the RHEB-Free complex. Nevertheless, all TM-scores exceeded the mean CASP14 and CASP15 benchmark values (both ∼0.91) across all categories (TBM-easy, TBM-hard, TBM/FM, and FM). The MDFF refinement protocol, prioritizes global fitting to cryo-EM density over optimizing fine-grained conformational details within individual subunits, which leads to a modest reduction in per-chain TM-scores relative to those for the FC. To further assess per-chain structural similarity to the experimental cryo-EM reference, we computed GDT-TS and GDT-HA scores and compared them against CASP benchmarks (ESI **Figure S5** and **Table S4**). For the RHEB-Bound system, average GDT-TS values for mTOR (0.72) and RHEB (0.79) chains were comparable to the mean scores of top-performing CASP13 models (0.70) across all categories, whereas chains corresponding to mLST8 (0.91) and RAPTOR (0.83) more closely matched the mean GDT-TS values observed in CASP14 and CASP15 (∼0.86). In the RHEB-Free system, mLST8 (0.90) and RAPTOR (0.88) exhibited GDT-TS values similar to those observed in the RHEB-Bound state, while the lowest GDT-TS values were observed for the mTOR chains (0.49), falling below the mean values for CASP13 TBM-hard and FM/TBM categories (∼0.60) but above the mean for CASP13 FM (∼0.40). Despite the relatively low score, the models for the RHEB-free mTOR chains still remain comparable to many of the highest-scoring CASP13 models within the TBM-hard, FM/TBM, and FM categories. For GDT-HA, the lowest scores were observed for the RHEB-unbound mTOR chains (0.28), yet these values remain comparable to several of the top-performing CASP13 models in the FM category, as illustrated in the lower region of the CASP13 violin plot (ESI **Figure S5** and **Table S4).** The next lowest GDT-HA values corresponded to the RHEB-bound mTOR chains (0.50), which still fall within the range of the best-performing CASP13 models across the TBM-hard, FM/TBM, and FM categories. In contrast, all other chains—including mLST8 and RAPTOR in both the RHEB-bound and RHEB-unbound complexes—exhibited GDT-HA values between 0.63 and 0.73, aligning well with those reported in CASP13–15 across all categories (**ESI Figure** S**5** and **Table S4**).

In addition to intra-chain validation, we assessed structural similarity of protein-protein interfaces (PPIs) by calculating Interface Similarity Scores (IS) and Interface TM-scores (ITM). IS- and ITM-scores for PPIs ranged from 0.30 to 0.73 and 0.42 to 0.82 respectively (**Figure 2C**), both correspond to highly significant p-values (<10⁻⁴ and <10⁻⁵, respectively) based on benchmarks (green dotted lines in **Figure 2**) against ∼1.8 million random interfaces (Gao et al., 2010). These values confirm that the modelled PPIs are statistically robust and closely recapitulate those observed in the experimental cryo-EM structures.

We noticed that all chain-specific structural scores exhibited a slight asymmetry for equilibrated models of the same protein copies in the two monomer halves of the complex (**Table S4**). To investigate whether this asymmetry increased during our longer production simulations, we compared the same set of scores for the protein copies in the two monomer halves of the complex at each local equilibria (plateaus in the 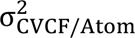 time trace) attained by the RHEB-Free and RHEB-Bound complex. Specifically, we extracted the average protein structure from trajectory segments of the complexes up to each plateau (vertical dashed lines in 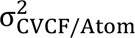 trace shown in ESI **Figure S6**). From these truncated trajectories, we identified and extracted all structures with the lowest RMSD relative to the corresponding average structure, thereby obtaining locally equilibrated structures associated with each 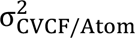 plateau. This procedure was repeated across multiple CVCF slope thresholds (10^-2^ − 10^-4^ Å^2^/ns) to define plateaus following the approach described in **STAR Methods (Complex Dynamics around the MD refined Cryo-EM State)**. The resulting scores (ESI **Figure S6**) were then compared with the mean values obtained for the two monomeric halves during the initial equilibration. Across all conditions, the scores remained close to values obtained at the end of the MD equilibration runs indicating that the simulations did not drift toward an unphysically or excessively asymmetric structural state.

To summarize, the refined models fully recapitulate structural features captured in the cryo-EM data (Yang et al., 2017) at local chain level, global complex level, and in terms of PPIs. Additionally, these refined models provide a high-resolution view of structural, dynamical and energetic changes induced in mTORC1 by RHEB as detailed in next subsections.

### High-Resolution MD Models of mTORC1 Reveal Global Conformational Changes and Allosteric Domain Shifts Induced by RHEB Binding

When examining the conformational changes within the Cryo-EM structures of mTORC1 in terms of relative displacements of protein chains or domains (**Figure 3A/C**), large error bars in atomic positions due to the limited resolution make it difficult to follow the changes induced by RHEB binding (**Figure 3B/D**, ESI **Figure S2-A, B** and **Figure S3-A, B)**. Starting from the AlphaFold predictions, the MD-equilibration and production runs refine the models to include realistic statistical distributions of centre of mass (COM) distances between the constituent proteins in the complex. Examining these distances across the multiple ±RHEB and ±ATP complexes, production trajectories and performing a statistical p-test we can identify significant distance changes between proteins in the complex induced by RHEB binding **Figure 3B** and ESI **Figure S2-A** show that in the +ATP state the mLST8 chains move away from mTOR as well as from each other after RHEB binding as evident from the changes in A-D, B-E and D-E COM distances. On the other hand, the RAPTOR protein chains come closer to the mTOR chains as well as each other when RHEB binds. Viewing the mTORC1 dimeric complex in **Figure 3A** as an ellipse, our MD trajectories show that RHEB binding significantly decreases the length of the major axis of the complex connecting the two RAPTOR copies and increases the length of the minor axis connecting the two mLST8 proteins (**Figure 3B**). Further, the distributions of the COM separations (violin plots in **Figure 3B**) show that the major axis is rigid with a single well-defined state for both ±RHEB complexes. On the other hand, the minor axis is flexible exhibiting multiple states for both complexes. These trends are slightly altered when ATP is not present at the mTOR active site. For instance, in the ATP-Free state the minor axis (D-E COM distance) shows no significant change. In contrast, a significant decrease in major axis length (W-Y COM distance) is still present.

**Figure 3:**
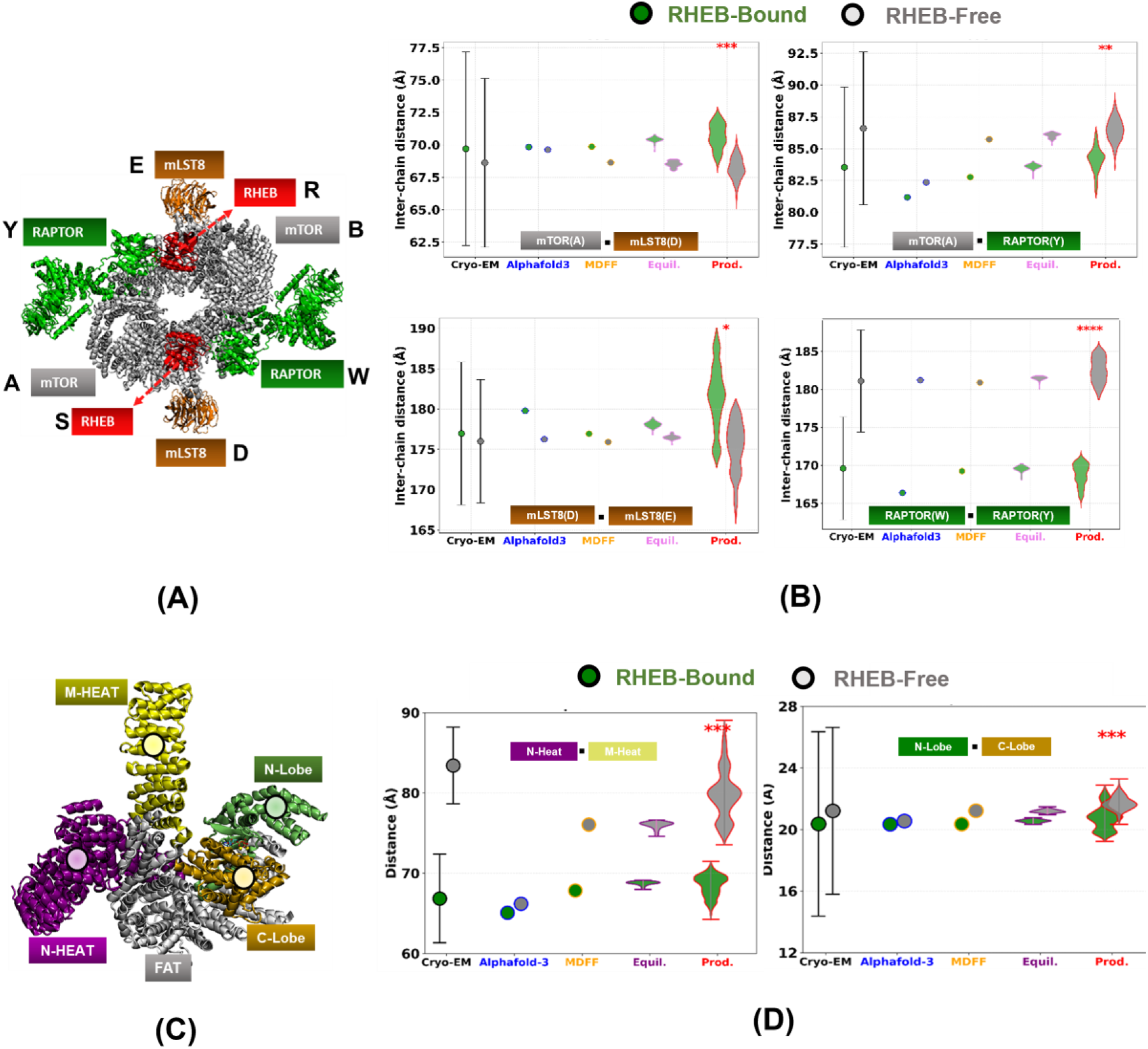
(A) Representative Cryo-EM Structure with RHEB bound (PDB ID: 6BCU) illustrating constituent protein chains with annotated chain ids (B) A comparison of the COM-COM separations of selected protein chains within complexes with and without RHEB from Cryo-EM experiments and models derived in this study (data for other protein chains are provided in ESI **Figure S2**). (C) Representative Cryo-EM structure of mTOR within the RHEB-Bound mTORC1 complex (6BCU). Different domains of the mTOR are coloured and labelled. (D) A comparison of the COM-COM separations of N-HEAT and M-HEAT domains (left) and Kinase Domain N-Lobe and C-Lobe for one pair of mTOR copies within RHEB-Bound and RHEB-free complexes (data for all combinations of mTOR copies are provided in ESI **Figure S3).** In panels B and D, we present data from the original Cryo-EM structures (6BCU and 6BCX), Alphafold3 based initial models, MDFF refined model, NPT-equilibrated ensemble, and NPT-production ensemble. Statistical significance (p-test) of changes in distances from multiple MD NPT-production runs are provided (ns(p > 0.05); * (p ≤ 0.05), **(p ≤ 0.01), ***p ≤ 0.001, ****p ≤ 0.0001.). Errors in COM distances for Cryo-EM structures were calculated via error propagation of atomic B-factors.

Cryo-EM studies have previously indicated that RHEB binding induces a major conformational shift in the mTORC1 complex both globally as well as in mTOR (**Figure 3D**), particularly involving compaction of the N-HEAT and M-HEAT domains(Yang et al., 2017) of mTOR. Specifically, a decrease in domain COM separations of N- and M-Heat domains of ∼16 Å can be computed from the reported structures. As shown in **Figure 3D** this change is well resolved in the Cryo-EM data. It has been further suggested that such changes lead to N-lobe and C-lobe domain movements in mTOR, which translate to a more compact kinase active site in the RHEB-free form of the complex (Chao and Avruch 2019). However, changes in the kinase N-/C-lobe separations induced by RHEB-binding (∼0.86 Å) are not resolved in the Cryo-EM data (**Figure 3D**). The non-uniform resolution across cryo-EM maps of the complexes, makes it difficult to verify such finer level structural changes. By contrast, our final MD equilibrated models offer improved spatial precision and interpretability. We observe a more consistent and tightly distributed N-/M-HEAT COM distance shift from 76.00 Å (-RHEB) to 68.77 Å (+RHEB), with standard deviations of only 0.24 Å and 0.44 Å, respectively (**Figure3D**). This refined measurement supports the hypothesis that RHEB binding brings N-HEAT and M-HEAT domains closer together, likely priming the kinase domain for catalysis. Further our refined MD models reveal a consistent 0.65 Å shift in the N-/C-lobe separations, which are well resolved over the thermal MD widths (±0.10 Å) (**Figure3(D)**). The resolution of this change is enhanced when we consider statistics from the multiple MD-production runs each of RHEB-Bound and RHEB-Free complexes. We observe a statistically significant (***) decrease in the N-C-lobe distance induced by RHEB binding. These results are summarized in **Figures 3B** and **3D** for one copy of mTOR from RHEB bound and RHEB unbound mTORC1 with ATP. Similar results were obtained with the same analysis for mTOR copies with different configurations (± ATP ± RHEB) as shown in ESI **Fig**ures **S3-4** and **Tables S2-S3**. We also observe the decrease in distance between N-/M-HEAT as well as N-/C-lobe upon RHEB binding also occurs in the absence of ATP. Taken together, these findings illustrate that the refined structural ensemble from MD not only recapitulates known chain or domain motions but also improves resolution and provides higher segment and residue level information within the kinase domain. As we see below these changes translate into both energetic and structural considerations at the active site,

### RHEB Induces a Favourable Enthalpy of ATP Binding to mTORC1

We carried out MM/GBSA enthalpy calculations for ATP binding to mTORC1 in the presence and absence of RHEB, as described in detail in **Methods (ATP Binding Energetics)**. The binding enthalpy (*ΔH*) calculations were performed for an active site of protein residues in mTOR lying within a sphere of variable cut-off radius (***r_cutoff_***) centred around the ATP with its two Mg^2+^ ions (**Figure 4A**). We find that independent of the cutoff radius, the presence of RHEB consistently produces more favourable binding enthalpy values (by ∼30 kcal/mol) relative to the RHEB-Free system (**Figure 4B** and ESI **Figures S7A**). We also see that this trend is already present in the gas-phase contributions alone (ESI **Figure S7C**) and preserved when polar solvent contributions are added (ESI **Figure S7E**). In fact, a *ΔH* decomposition (ESI **Table S5**) reveals that non-bonded electrostatic contributions dominate the ATP binding enthalpy for both ± RHEB systems. RHEB binding induces more favourable electrostatics with an enthalpy stabilization of ∼100-180 kcal/mol, which increases with ***r_cutoff_***. On the other hand, the much weaker van der Waals contributions tend to become more unfavourable (by ∼8-22 kcal/mol) with RHEB binding. To determine whether the more favourable *ΔH* for ATP binding in the presence of RHEB arises from strengthened interactions between ligand and mTOR active-site residues, we computed the nonbonded interaction energies between the former and later as a function of ***r_cutoff_*** for the two ± RHEB variants of the ATP bound complex. Surprisingly, these calculations revealed that independent of ***r_cutoff_***, ATP–protein non-bonded interactions are counterintuitively stronger in the RHEB-Free system (**Figure 4C** and ESI **Figure S7B**). This implies that the enhanced binding enthalpy observed in the RHEB-bound complex does not originate from more favourable direct ATP–active site interactions. Rather, RHEB binding induces an active site pre-organization creating an energetically favourable active site of residues surrounding the ATP + 2Mg²⁺, thereby contributing to the overall enthalpic stabilization.

**Figure 4:**
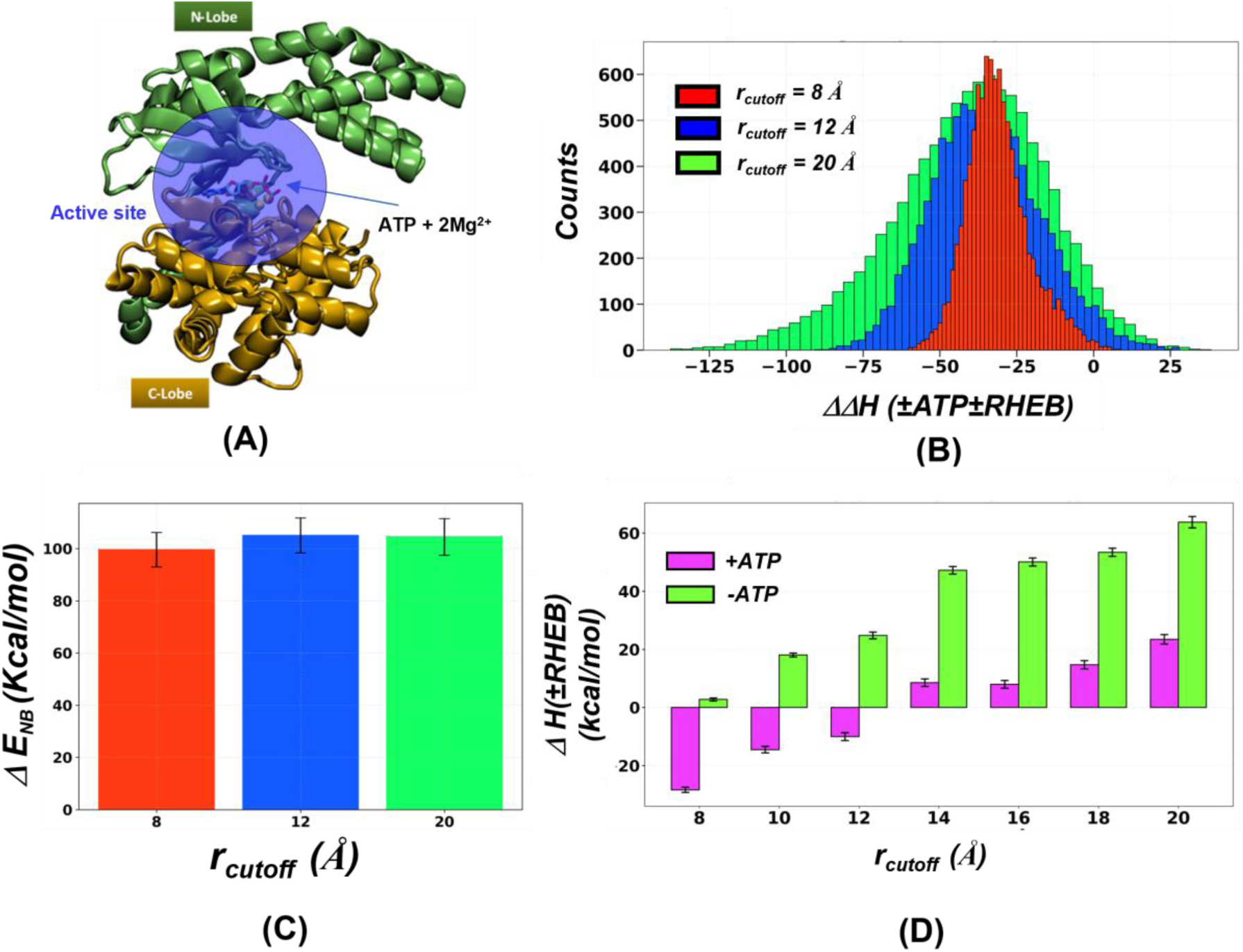
(A) ATP+2Mg^2+^ docked between N-/C-Lobe of the mTOR KD. All energy calculations are carried out for an active site of residues lying within a sphere of radius **r_cutoff_** centred at the COM of ATP+2Mg^2+^ (B) Distributions of the change in enthalpy of ATP binding (**Eqn.11**) induced by presence of RHEB ΔΔH(±ATP±RHEB) (C) Change in non-bonded interaction energy (Δ**E_NB_**) between ATP+2Mg^2+^ and surrounding residues induced by RHEB binding. Distributions of ΔΔH (10^4^ values) and Δ**E_NB_** (10^2^ values) are calculated as a function of ***r_cutoff_*** from NPT-production trajectories for mTORC1 ± 2 × ATP ± 2 × RHEB (20 trajectories) and mTORC1 + 2 × ATP ± 2 × RHEB (10 trajectories) respectively. (D) Change in the enthalpy of mTORC1 (ΔH(±RHEB)) induced by RHEB binding in the presence and absence of ATP.

We next investigated whether RHEB influences the +ATP and/or the -ATP states of mTORC1 to make ATP binding more favourable. For this we computed the change in enthalpy with RHEB binding (*ΔH*(±RHEB)) as a function of ***r_cutoff_*** for the +ATP and -ATP states of the complex separately (**Figure 4D**). Interestingly, RHEB binding impacts the +ATP and -ATP states in very different ways, which are sensitive to the ***r_cutoff_*** values. The -ATP state is destabilized enthalpically to increasing extents (∼2-62 kcal/mol) as ***r_cutoff_*** is increased from 8-20 Å. On the other hand, the +ATP state is initially enthalpically stabilized at small values of ***r_cutoff_*** (8-12Å), but gets progressively destabilized at larger cutoff values (14-20 Å). While the RHEB induced stabilization of the +ATP complex dominates over the destabilization of the - ATP complex close to the binding site (***r_cutoff_*** = 8 Å) the latter dominates over longer ranges. Overall, the differential impact of RHEB on the +ATP and -ATP forms leads to a favourable ATP-binding enthalpy which is insensitive to the active site cut-off as seen in **Figure 4B** and ESI **Figure S7A**.

### RHEB Activation Alters ATP Conformation and Interactions in the KD

The altered enthalpy of the +ATP state of the complex in the presence of RHEB indicates that the ATP docking conformation and the interactions at the active site are altered by the effector. We therefore examined the solvent accessibility of the ATP, its interactions with the catalytic D2338, which acts as a base during substrate phosphorylation (Yang et al., 2013b), and the coordination sphere of the Mg^2+^ in the presence and absence of RHEB. We find that the solvent accessibility of ATP lies in the range of 10-35 % in the MD structural ensembles from the production runs (**Figure 5(A)** and ESI **Figure S8**) with two distinct pSASA_ATP_ peaks in the RHEB-Bound form and a single broad peak for the RHEB-Free form. In the absence of RHEB, ATP remains deeply buried exhibiting a peak pSASA_ATP_ of ∼19 %. However, RHEB binding creates two distinct populations of ATP conformations, one with a low solvent accessibility (∼16 %) and the other with a higher exposure to solvent (∼27%). Upon analysing the critical catalytic Asp2338(γC)-ATP(γP) distance across all trajectories, we again observed a bimodal distribution of shorter distances (peaks at ∼10 Å and ∼8.5 Å) in RHEB-Bound and unimodal distribution of longer distances (peak at ∼11.5 Å) in RHEB-Free trajectories (shaded distributions in **Figure 5B**). To correlate the Asp2338(γC)-ATP(γP) distances with the three subpopulations of ATP conformations with different solvent exposures in the presence and absence of RHEB, we extracted the structural ensembles of ATP, Mg^2+^ and surrounding active site residues (***r_cutoff_*** = 3Å) for the buried and solvent exposed ATP populations in RHEB-Bound (*C_BB_* and *C_EB_*) and the buried ATP population in RHEB-Free (*C_BF_*) trajectories (**Figure 5A**). The D2338(γC)-ATP(γP) distances extracted from these three subpopulations (box plots in **Figure 5B**) perfectly match with the peaks of distance distributions across MD trajectories. This indicates that the two conformations of ATP in the presence of RHEB (*C_BB_* and *C_EB_*) are positioned closer to D2338. This observation gains significance in the context of a recent Cryo-EM study which shows RHEB binding to mTORC1 attached to lysosomal membranes leads to a more potent (active) complex (Cui et al., 2025). As seen in **Figure 5B** D2338(γC)-ATP(γP) distances in the complex are shorter when RHEB is bound and more consistent with those from the “intermediate” or “fully activated” complexes discussed by Hurley and co-workers (Cui et al., 2025). We note that conformational preorganization upon substrate docking can further decrease the D2338(γC)-ATP(γP) distances seen in the +RHEB complex to make the active site more catalytically competent. Nevertheless, even without the substrate, the mean D2338(γC)-ATP(γP) distances for the *C_EB_* ensemble (∼8.4Å) are comparable to analogous interactions in other systems like the ERK2 kinase or insulin receptor kinase (ESI **Table S6**).

**Figure 5:**
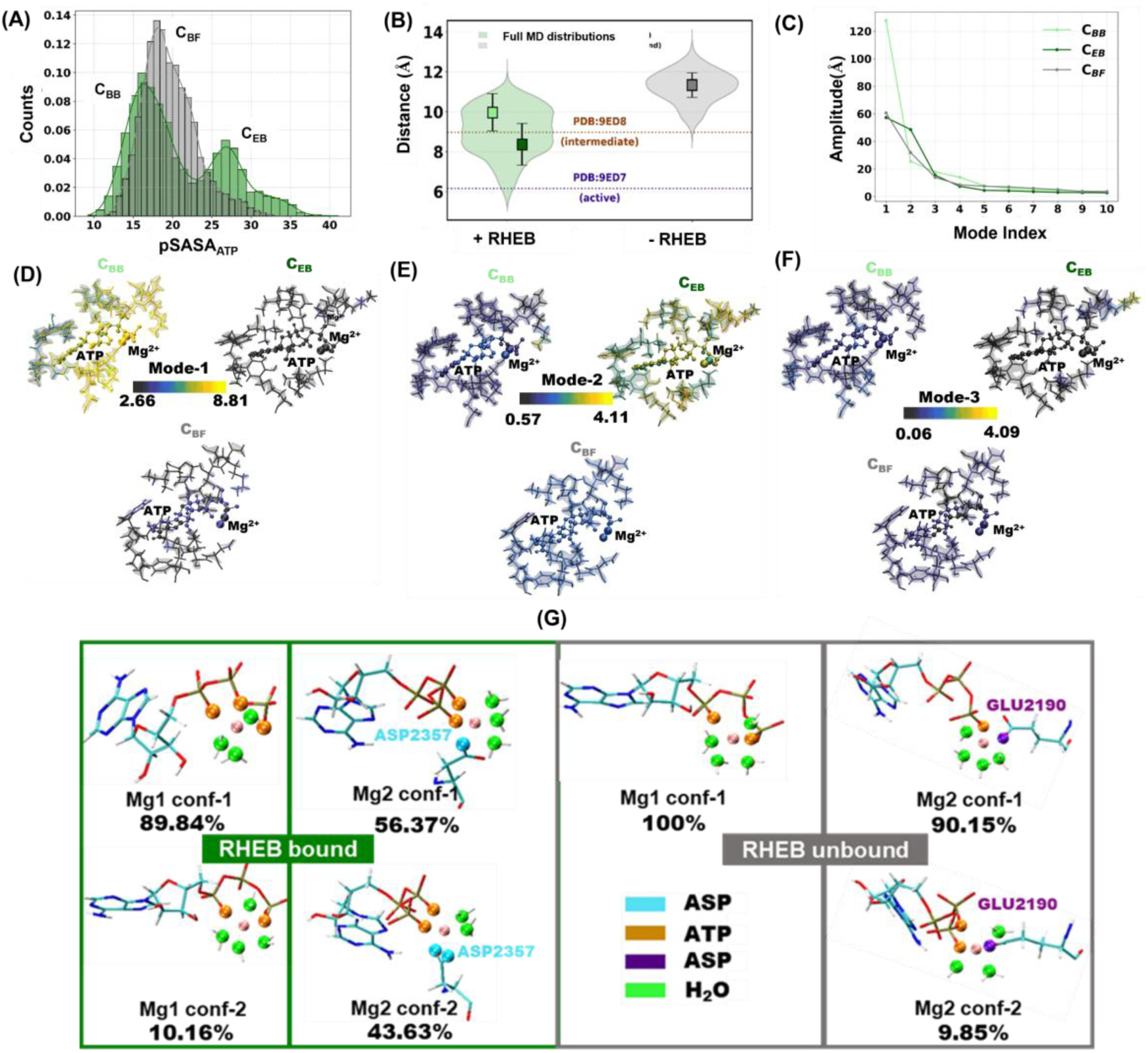
(A) Distributions of ATP solvent accessibility from NPT-production trajectories (last 5 ns from each instance). (B) Box plots for the distance between D2338 and ATP-γP for RHEB-Bound **C_BB_**/**C_EB_** (buried/exposed ATP) ensembles in dark/light green and RHEB-Free **C_FB_** (buried ATP) ensemble in dark grey. The shaded violin plots show distance distributions from the full NPT-trajectories. The horizontal dotted lines indicate values from recent Cryo-EM structures of reconstituted mTORC1–RHEB–RAG–Ragulator with the full-length substrate 4E-BP1 complex on liposomes (C) Mode Amplitude spectrum. (D-F) Top 3 PC modes of active site structural ensembles for the three ATP subpopulations in the RHEB-Bound and RHEB-Free MD trajectories. (G) Coordination modes of Mg1 and Mg2 in RHEB-Bound(left) and RHEB-Free(right) complexes with their relative abundance in MD production trajectories.

Note that the D2338(γC)-ATP(γP) distances in the original mTORC1 complexes before MD refinement show insignificant differences (∼0.5 Å) with and without RHEB (**TableS6**). Interestingly, the buried ATP in the *C_BB_* state shows substantially greater conformational heterogeneity than in the *C_BF_* state or even in the more solvent exposed *C_EB_* state as seen from the amplitude spectrum of PCs captured for these ensembles (**Figure 5C**). The top PC mode reveals much larger conformational fluctuations for the *C_BB_* state relative to the other two ATP ensembles, particularly among residues surrounding the γ-phosphate of ATP and the coordinating Mg²⁺ ions regions corresponding to the catalytic cleft (**Figure 5D**). PC mode 2 shows enhanced fluctuations for the more solvent exposed *C_EB_* state relative to the other two states (**Figure 5E**). Finally, higher modes show much lower fluctuations and do not significantly distinguish between the three different ATP states in the complexes.

The coordination spheres of the two Mg^2+^ ions (Mg1 and Mg2) appear to be also influenced by RHEB binding as revealed from our simulation ensemble—ten replicates for each monomeric unit in the +RHEB and -RHEB states. As shown in the **Figure 5G**, Mg1 in the +RHEB complex adopts two distinct coordination patterns: a predominant mode (*conf-1*) in which three ATP tail oxygens and three water molecules serve as ligands, and a less populated mode (*conf-2*) in which two ATP oxygens and four waters completed the coordination sphere. In the -RHEB form, Mg1 exhibits only the latter coordination pattern (*conf-2*). In contrast, Mg2 displays marked differences in its coordination sphere in the presence and absence of RHEB. While two coordination modes are observed for both states (**Figure 5G**), a key distinction induced by RHEB binding is the replacement of the glutamate side-chain ligand observed in cryo-EM by an aspartate (D2357), whose side-chain carboxylate can donate either one (*conf-1*) or two coordinating oxygens (*conf-2*) to Mg2. Furthermore, RHEB binding reduces the number of water ligands and increases ATP ligation for both Mg²⁺ ions which may play a role in positioning the molecule in relation to the incoming substrate.

Taken together, these results indicate that RHEB binding induces ATP states with increased conformational dynamics in the catalytic cleft and favourable interactions with the active site aspartates and Mg^2+^ ions. These changes can be linked to substrate accommodation and ATP positioning during catalysis. —changes that may contribute directly to RHEB-mediated activation of the mTOR kinase.

### RHEB Stabilizes the mTORC1 Complex and Allosterically Alters the Dynamics of Kinase and FAT Domains

To investigate the dynamic impact of RHEB binding on mTORC1, we examined time traces of the cumulative variance (σ*^2^_CVCF_*(𝑇_𝑠_)) for different subsets of atoms (**Figure 6** and ESI **Figure S11**): the structured backbone *C_α_* atoms of the entire complex and individual chains, and specific domains within mTOR (N-HEAT,M-HEAT,FAT,KD-N-Lobe, and KD-C-Lobe). The trace of the cumulative variance (σ*^2^_CVCF_*) of atomic fluctuations along MD trajectories tracks the features of the underlying protein energy landscape (Paul et al., 2020) (Das & Venkatramani, 2025). A plateau in the σ*^2^_CVCF_* trace indicates local equilibration for a section of the landscape, whereas an increasing variance indicates progressive conformational exploration. At local equilibria the values of σ*^2^_CVCF_* can be formally connected to the curvature of the underlying energy landscape by the equipartion theorem helping distinguish flexibility of systems. (Paul et. al. 2021)

**Figure 6:**
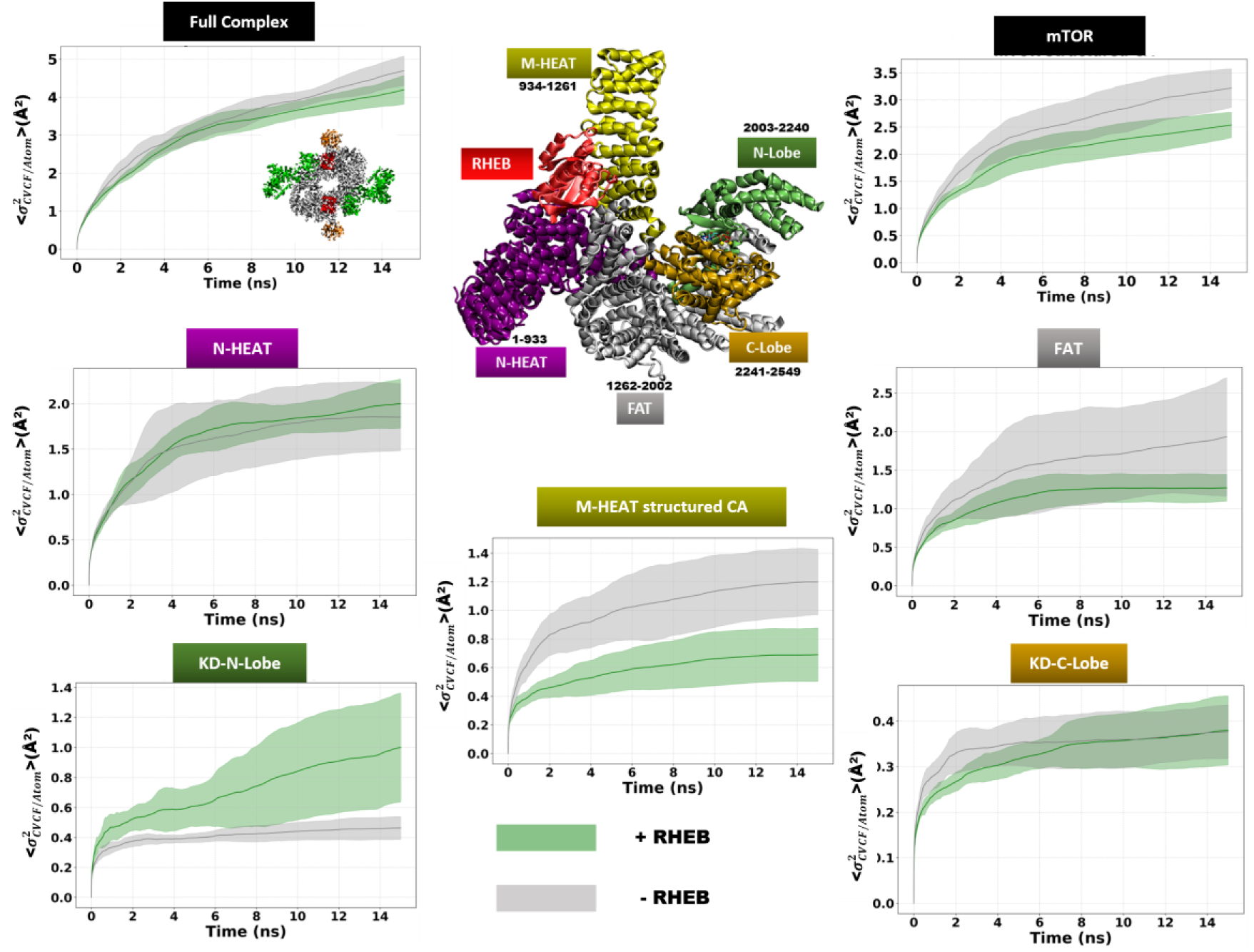
Mean and std.dev of per atom σ*^2^_CVCF_* traces of structured *C* atoms of Full Complex (top left) along with mTOR (top right) along with plots for the same for respective domains constituting mTOR with ATP from 5 × 15ns MD trajectories for full mTORC1 and 10 × 15ns (two instances for each complex) for mTOR chains and respective domains with (green) and without (grey) RHEB. The inset structure in the top row shows the mTOR along with its constituent domains and their interactions with RHEB.

The time trace of the σ*^2^_CVCF_* across 5 trajectories for the full complex (**Figure 6** and ESI **Figure S11-13**) reveals that all four ±RHEB ±ATP mTORC1 complexes do not fully locally achieve equilibrium sampling (on average) over the timescale of 15 ns. Investigation of the raw traces indicate that RHEB tends to stabilize the complex producing more stable sections of the trajectories (ESI **Figures S11** and **S13**). While the overlap of the traces indicates that the changes in dynamics with RHEB binding is not fully resolved for the overall mTORC1 complex, the variance of the individual chains (**Figure 6** and ESI **Figures S11-S16**) for ±ATP systems exhibit diversity. It is clear that the biggest contribution to the dynamics of the complex arises from the mTOR protein chains with smaller contributions from the RAPTOR units and negligible contribution from mLST8 (ESI **Figure S16**). The σ*^2^_CVCF_* traces indicate that in both ±ATP states RHEB binding decreases the dynamics for mTOR and to a lesser extent in mLST8 but does not alter that of the RAPTOR chains. Within mTOR there is a clear asymmetry in the response of different domains to RHEB binding. The clearest signatures are seen for the kinase N-lobe, FAT and the M-HEAT domains which register opposing changes. RHEB binding stabilizes the FAT domain and increases the dynamics of the kinase N-lobe, specifically destabilizing it, which correlates with increased dynamics in the kinase catalytic cleft observed in previous sections. On the other hand, the kinase C-lobe is not impacted in terms of dynamics by the presence of RHEB. These trends are independent of whether ATP is present at the kinase active site. The changes in N-lobe, and FAT domain dynamics induced by RHEB are allosteric in nature as the activator only directly interfaces with the M-HEAT and N-HEAT domains (**Figure 6** inset structure). As known previously, and refined by our models here, RHEB binding brings these two domains closer together and significantly narrows the distribution of M/N-Heat separations (**Figure 3D**). Here, our σ*^2^_CVCF_* trace analysis further reveals that RHEB binding significantly reduces the flexibility of the M-HEAT domain in both ±ATP states while increasing the flexibility of the N-HEAT domain in the -ATP state selectively. The effect of RHEB on dynamics of the complex, constituent protein chains, and mTOR domains which are already apparent in 15 ns trajectories are even more clearly established when the sets of +ATP trajectories of the ±RHEB complexes were extended up to 25 ns (ESI **Figures S14-S15** and **S17**).

## Discussion

The Cryo-EM study by Pavletich group (Yang et al., 2017) provided the first structural basis to understanding the allosteric activation of mTORC1 by RHEB. However, the varied resolution across the complex in the Cryo-EM structures where 16% of residues are missing, make it difficult to track high resolution structural changes induced by RHEB binding. Further, the structures cannot be directly utilized to carry out dynamical analysis using techniques such as MD simulations. Here, by combining the predictive power of AlphaFold-3 with steered molecular dynamics (MDFF) and a multi-stage equilibration protocol, we bridge the gap between Cryo-EM data and high-resolution molecular models suitable for dynamic analyses.

Over the past half a decade, AlphaFold has emerged as a powerful AI tool to reliably predict structures of proteins from their primary sequence. The latest AlphaFold-3 algorithm extends this capability to protein complexes, enabling the prediction of protein–protein contacts, flexible loop positioning, and even cofactor placement. The large size of the mTORc1 dimeric complex and the large-scale conformation changes induced in it by RHEB binding preclude the direct application of AlphaFold to this system. Thus, we modelled each monomeric unit within the complexes using AlphaFold and aligned them to the Cryo-EM structures to generate initial models of mTORC1 with and without RHEB. These initial models do not capture RHEB induced large scale structural changes observed in the Cryo-EM and further refinement was necessary. The use of cryo-EM densities as structural restraints in MDFF simulations of AlphaFold-based initial models created structurally complete models, which capture critical large scale allosteric shifts captured in experiments (**Figures 2** and **3D**). Following this, we applied a progressive MD equilibration scheme to selectively relax experimentally resolved and unresolved regions through gradually decreasing restraints. This strategy preserves the fidelity of cryo-EM-fitted regions while allowing flexible parts of the structure to settle into low-energy conformations. The result is a kinetically equilibrated thermal ensemble of complex geometries that also reflects the experimentally resolved structural features of mTORC1 in presence and absence of RHEB (**Figure 3D**) as validated by local, global, and interfacial structural metrics (**Figure 2**). At the same time, the refined model provides a dynamical high-resolution view of the ATP coordination at the active site as well as local and global conformational rearrangements. Further, we have carried out short MD-production runs for the complex to assess the energy landscape and resulting dynamics around the refined model geometries. Using these dynamically robust models, we find RHEB induced structural and dynamical changes at FC, domain, and kinase active site levels.

On the scale of the ellipsoidal complex RHEB binding decreases the length of the rigid major axis connecting the two RAPTOR copies which leads to a tighter mTOR-Raptor packing. On the other hand, the flexible minor axis connecting the mLST8 copies is increased upon RHEB binding loosening mTOR-mLST8 interactions (**Figure 3B** and ESI **Figure S2**). Our refined models retain the reduction in N-Heat and M-Heat domain separation induced by RHEB. However, the reduction is predicted to be more modest (∼7 Å) relative to that observed in the original Cryo-EM structures (∼16 Å). At a finer level our models verify that these domain level conformational changes are translated to the kinase active site. The kinase N-lobe and C-lobe are reoriented (**Figure 3D**) such that the active site is enthalpically more favourable for ATP-binding. MM-GBSA calculations on the sets of NPT-production trajectories for all four complexes (mTORC1 ± RHEB ± ATP) reveal that the presence of RHEB makes ATP binding enthalpically more favourable by ∼25 kcal/mol (**Figure 4B**). RHEB achieves this allosterically by simultaneously stabilizing the active site electrostatic interactions in +ATP state and destabilizing that in the -ATP state of the kinase. Our analysis (**Figure 4D**) further reveals that RHEB impacts the +ATP state more in the vicinity of the ligand and the -ATP state further away (≥ 10Å). Surprisingly, the long-range structural changes in mTORC1 brought about by RHEB binding promote more favourable protein–ligand interactions, despite weakening direct nonbonded interactions of the ATP with active-site residues (**Figure 4B**). Our simulations further reveal that RHEB changes the active-site conformations of ATP creating two distinct subpopulations (*C_BB_* and *C_EB_*) both of which are better positioned for catalysis relative to the single population (*C_BF_*) observed in the -RHEB models (**Figure 5A-B**). Specifically, RHEB increases the proximity of the ATP tail to D2338 implicated in substrate phosphorylation (Yang et al., 2013b) pushing the complex towards an activated state (Cui et al., 2025). RHEB binding also remodels the coordination environment of both Mg²⁺ ions, replacing water coordination with that from the ATP phosphate tail substituting of Glu-based coordination with Asp-based coordination.

mTORC1 dynamics as extracted from MD production trajectories also indicates that the presence of RHEB is favourable to achieve a catalytically competent state. For instance, the less solvent exposed ATP state in the +RHEB complex (*C_BB_*) displays markedly higher conformational fluctuations near the catalytic cleft than the other states in the presence and absence of RHEB (**Figure 5C**), a feature that may facilitate improved catalytic efficiency through enhanced substrate accommodation. Our dynamical (σ*^2^_CVCF_* trace) analysis of all four mTORC1 ± RHEB ± ATP complexes (**Figure 6** and ESI **Figure S11-S17**) reveals that the increased thermal fluctuations at the kinase catalytic site originate from a larger allosteric destabilization of the N-lobe by RHEB. Interestingly, RHEB also allosterically stabilizes the kinase FAT domain and leaves the dynamics of the C-lobe unchanged. Further, and more directly RHEB interfaces with the M-HEAT and N-HEAT domains bringing them closer and at the same time reducing their dynamics by stabilizing the M-HEAT domain. Overall, these direct and allosteric dynamical changes brought about by RHEB lead to a stabilization of the mTOR chain and at a more global scale, the mTORc1 complex itself. RHEB binding thus potentially possesses an additional mechanistic lever in the form of dynamics for tuning the rate of phosphoryl transfer by the complex. A more complete dynamical study of the multiple complex states (mTORC1 ± RHEB ± ATP ± Substrate) with better sampling statistics would be required to fully understand the reshaping of the kinase energy landscape for catalysis by RHEB. The present study sets up refined solvated models for such efforts which are presently underway in our group.

## Conclusion

Cryo-EM studies have provided unprecedented access to the functionally relevant states of large complexes. However, non-uniform resolution of constituent protein chains presents a significant barrier towards developing quantitative insights. Here, we establish a comprehensive pipeline that unifies AlphaFold-3 prediction, MDFF refinement, and progressive equilibration to generate chemically accurate and dynamically stable models of large multi-protein complexes. Refined models of mTORC1 built from existing Cryo-EM data reveal how RHEB binding drives global architectural rearrangements, enhances ATP binding energetics, reshapes Mg²⁺ coordination, and allosterically activates the kinase domain. Our results highlight how allosteric activation emerges from the interplay between global structural and dynamical changes which translate into a catalytically competent active-site reorganization and dynamics, offering a high-resolution mechanistic understanding of RHEB-mediated mTORC1 activation. The resulting models, together with our modelling pipeline, offer a versatile platform for probing diverse regulatory influences on mTORC1 such as the binding of protein activators/inhibitors, mutations, and post translational modifications.

## Lead Author

Requests for further information and resources should be directed to and will be fulfilled by the lead contact, Prof. Ravindra Venkatramani.

## Acknowledgement

The authors acknowledge the funding support from the Department of Atomic Energy (DAE), Government of India, under Project Identification No. 1303/10/2025-R&D-II-DAE/TIFR-17248 RTI 4015. PG acknowledges Dr. Krishnakant Vishwakarma, Dr. Anustup Chakraborty, and Dr. Mitradip Das for valuable scientific discussions and conceptual input on dynamical analysis, MMGBSA calculations and quasi-harmonic analysis.

## References

Abramson, J., Adler, J., Dunger, J., Evans, R., Green, T., Pritzel, A., Ronneberger, O., Willmore, L., Ballard, A. J., Bambrick, J., Bodenstein, S. W., Evans, D. A., Hung, C. C., O’Neill, M., Reiman, D., Tunyasuvunakool, K., Wu, Z., Žemgulytė, A., Arvaniti, E., … Jumper, J. M. (2024). Accurate structure prediction of biomolecular interactions with AlphaFold 3. Nature 2024 630:8016, 630(8016), 493–500. 10.1038/s41586-024-07487-w

Ali, E. S., Mitra, K., Akter, S., Ramproshad, S., Mondal, B., Khan, I. N., Islam, M. T., Sharifi-Rad, J., Calina, D., & Cho, W. C. (2022). Recent advances and limitations of mTOR inhibitors in the treatment of cancer. Cancer Cell International 2022 22:1, 22(1), 1–16. 10.1186/S12935-022-02706-8

Aylett, C. H. S., Sauer, E., Imseng, S., Boehringer, D., Hall, M. N., Ban, N., & Maier, T. (2016). Architecture of human mTOR complex 1. Science, 351(6268), 48–52. 10.1126/SCIENCE.AAA3870/SUPPL_FILE/AYLETT.SM.PDF

Biasini, M., Schmidt, T., Bienert, S., Mariani, V., Studer, G., Haas, J., Johner, N., Schenk, A. D., Philippsen, A., & Schwede, T. (2013). OpenStructure: An integrated software framework for computational structural biology. Acta Crystallographica Section D: Biological Crystallography, 69(5), 701–709. 10.1107/S0907444913007051/IC5090SUP2.TXT

Callaway, E. (2020). “It will change everything”: DeepMind’s AI makes gigantic leap in solving protein structures. Nature, 588(7837), 203–204. 10.1038/D41586-020-03348-4;SUBJMETA

Cui, Z., Esposito, A., Napolitano, G., Ballabio, A., & Hurley, J. H. (2025). Structural basis for mTORC1 activation on the lysosomal membrane. Nature, 647(8089), 536–543. 10.1038/S41586-025-09545-3;TECHMETA

Das, M., & Venkatramani, R. (2025). AutoSIM: Redesigning and automating umbrella sampling for biomolecular conformational transitions. Journal of Chemical Physics, 163(1), 14111. 10.1063/5.0268626

Gao, M., Skolnick, J., & Rost, B. (2010). iAlign: a method for the structural comparison of protein– protein interfaces. Bioinformatics, 26(18), 2259–2265. 10.1093/BIOINFORMATICS/BTQ404

Genheden, S., & Ryde, U. (2015). The MM/PBSA and MM/GBSA methods to estimate ligand-binding affinities. Expert Opinion on Drug Discovery, 10(5), 449. 10.1517/17460441.2015.1032936

Grabiner, B. C., Nardi, V., Birsoy, K., Possemato, R., Shen, K., Sinha, S., Jordan, A., Beck, A. H., & Sabatini, D. M. (2014). A diverse array of cancer-associated MTOR mutations are hyperactivating and can predict rapamycin sensitivity. Cancer Discovery, 4(5), 554–563. 10.1158/2159-8290.CD-13-0929/42520/AM/A-DIVERSE-ARRAY-OF-CANCER-ASSOCIATED-MTOR

Guzenko, D., Lafita, A., Monastyrskyy, B., Kryshtafovych, A., & Duarte, J. M. (2019). Assessment of protein assembly prediction in CASP13. Proteins: Structure, Function, and Bioinformatics, 87(12), 1190–1199. 10.1002/PROT.25795

Huang, J., & Mackerell, A. D. (2013). CHARMM36 all-atom additive protein force field: validation based on comparison to NMR data. Journal of Computational Chemistry, 34(25), 2135–2145. 10.1002/JCC.23354

Humphrey, W., Dalke, A., & Schulten, K. (1996). VMD: Visual molecular dynamics. Journal of Molecular Graphics, 14(1), 33–38. 10.1016/0263-7855(96)00018-5

Jumper, J., Evans, R., Pritzel, A., Green, T., Figurnov, M., Ronneberger, O., Tunyasuvunakool, K., Bates, R., Žídek, A., Potapenko, A., Bridgland, A., Meyer, C., Kohl, S. A. A., Ballard, A. J., Cowie, A., Romera-Paredes, B., Nikolov, S., Jain, R., Adler, J., … Hassabis, D. (2021). Highly accurate protein structure prediction with AlphaFold. Nature 2021 596:7873, 596(7873), 583–589. 10.1038/s41586-021-03819-2

Mariani, V., Biasini, M., Barbato, A., & Schwede, T. (2013). lDDT: a local superposition-free score for comparing protein structures and models using distance difference tests. Bioinformatics, 29(21), 2722. 10.1093/BIOINFORMATICS/BTT473

Mukherjee, S., & Zhang, Y. (2009). MM-align: a quick algorithm for aligning multiple-chain protein complex structures using iterative dynamic programming. Nucleic Acids Research, 37(11), e83– e83. 10.1093/NAR/GKP318

Ozden, B., Kryshtafovych, A., & Karaca, E. (2021). Assessment of the CASP14 assembly predictions. Proteins: Structure, Function, and Bioinformatics, 89(12), 1787–1799. 10.1002/PROT.26199

Ozden, B., Kryshtafovych, A., & Karaca, E. (2023). The impact of AI-based modeling on the accuracy of protein assembly prediction: Insights from CASP15. Proteins: Structure, Function, and Bioinformatics, 91(12), 1636–1657. 10.1002/PROT.26598

Paul, S., Ainavarapu, S. R. K., & Venkatramani, R. (2020). Variance of Atomic Coordinates as a Dynamical Metric to Distinguish Proteins and Protein-Protein Interactions in Molecular Dynamics Simulations. Journal of Physical Chemistry B, 124(21), 4247–4262. 10.1021/ACS.JPCB.0C01191/ASSET/IMAGES/LARGE/JP0C01191_0006.JPEG

Paul, S., Audhya, A., & Cui, Q. (2023). Molecular mechanism of GTP binding- and dimerization-induced enhancement of Sar1-mediated membrane remodeling. Proceedings of the National Academy of Sciences of the United States of America, 120(8), e2212513120. 10.1073/PNAS.2212513120;PAGE:STRING:ARTICLE/CHAPTER

Phillips, J. C., Hardy, D. J., Maia, J. D. C., Stone, J. E., Ribeiro, J. V., Bernardi, R. C., Buch, R., Fiorin, G., Hénin, J., Jiang, W., McGreevy, R., Melo, M. C. R., Radak, B. K., Skeel, R. D., Singharoy, A., Wang, Y., Roux, B., Aksimentiev, A., Luthey-Schulten, Z., … Tajkhorshid, E. (2020). Scalable molecular dynamics on CPU and GPU architectures with NAMD. Journal of Chemical Physics, 153(4), 44130. 10.1063/5.0014475/16709547/044130_1_ACCEPTED_MANUSCRIPT.PDF

Sancak, Y., Thoreen, C. C., Peterson, T. R., Lindquist, R. A., Kang, S. A., Spooner, E., Carr, S. A., & Sabatini, D. M. (2007). PRAS40 is an insulin-regulated inhibitor of the mTORC1 protein kinase. Molecular Cell, 25(6), 903–915. 10.1016/J.MOLCEL.2007.03.003

Shams, R., Ito, Y., & Miyatake, H. (2022). Development of an RHEB-Targeting Peptide to Inhibit mTORC1 Kinase Activity. ACS Omega, 7(27), 23479–23486. 10.1021/ACSOMEGA.2C01865/ASSET/IMAGES/MEDIUM/AO2C01865_M002.GIF

Tafur, L., Kefauver, J., Loewith, R., Ch, L. T., Kefauver@unige, J., & Ch, J. K. (2020). Structural Insights into TOR Signaling. Genes 2020, Vol. 11, Page 885, 11(8), 885. 10.3390/GENES11080885

Trabuco, L. G., Villa, E., Mitra, K., Frank, J., & Schulten, K. (2008). Flexible Fitting of Atomic Structures into Electron Microscopy Maps Using Molecular Dynamics. *Structure (London*, England : 1993), *16*(5), 673. 10.1016/J.STR.2008.03.005

Trabuco, L. G., Villa, E., Schreiner, E., Harrison, C. B., & Schulten, K. (2009). Molecular dynamics flexible fitting: a practical guide to combine cryo-electron microscopy and X-ray crystallography. *Methods (San Diego*, Calif.), 49(2), 174–180. 10.1016/J.YMETH.2009.04.005

Wälchli, M., Berneiser, K., Mangia, F., Imseng, S., Craigie, L. M., Stuttfeld, E., Hall, M. N., & Maier, T. (2021). Regulation of human mTOR complexes by DEPTOR. ELife, 10. 10.7554/ELIFE.70871

Woo, H., Park, S. J., Choi, Y. K., Park, T., Tanveer, M., Cao, Y., Kern, N. R., Lee, J., Yeom, M. S., Croll, T. I., Seok, C., & Im, W. (2020). Developing a fully glycosylated full-length SARS-COV-2 spike protein model in a viral membrane. Journal of Physical Chemistry B, 124(33), 7128– 7137. 10.1021/ACS.JPCB.0C04553/SUPPL_FILE/JP0C04553_SI_001.MP4

Yang, H., Jiang, X., Li, B., Yang, H. J., Miller, M., Yang, A., Dhar, A., & Pavletich, N. P. (2017). Mechanisms of mTORC1 activation by RHEB and inhibition by PRAS40. Nature 2017 552:7685, 552(7685), 368–373. 10.1038/nature25023

Yang, H., Rudge, D. G., Koos, J. D., Vaidialingam, B., Yang, H. J., & Pavletich, N. P. (2013a). mTOR kinase structure, mechanism and regulation. Nature 2013 497:7448, 497(7448), 217–223. 10.1038/nature12122

Yang, H., Rudge, D. G., Koos, J. D., Vaidialingam, B., Yang, H. J., & Pavletich, N. P. (2013b). mTOR kinase structure, mechanism and regulation. Nature 2013 497:7448, 497(7448), 217–223. 10.1038/nature12122

Zemla, A. (2003). LGA: a method for finding 3D similarities in protein structures. Nucleic Acids Research, 31(13), 3370. 10.1093/NAR/GKG571

Zhang, Y., & Skolnick, J. (2005). TM-align: a protein structure alignment algorithm based on the TM-score. Nucleic Acids Research, 33(7), 2302–2309. 10.1093/NAR/GKI524

