## Supporting Information Tables and Figures for "High Resolution Solvated Models Reveal Mechanisms of Allosteric Activation of mTORC1 by RHEB"

### Supplementary Information

### Table of Contents (Table)


| Item | Descrption | Page no. |
|---|---|---|
| **Table S1** | *Details of Solvated models used in Molecular Dynamics Simulations.* | **3** |
| **Table S2-A,B** | *N-/M-HEAT COM distance during Equilibration with, without ATP+2$Mg^{2+}$* | **9-10** |
| **Table S3-A,B** | *N-/C-Lobe COM distance during Equilibration with, without ATP+2$Mg^{2+}$* | **11-12** |
| **Table S4** | *Table with model validation metrics scores with errors.* | **13** |
| **Table S5** | *Individual Energy contributions to the Enthalpy change for RHEB bound(green) and RHEB unbound (grey).* | **17** |
| **Table S6** | *Distance between catalytic Asp with γ-Phosphate/mimic* | **28** |


### Table of Contents (Figure)


| Item | Descrption | Page no. |
|---|---|---|
| **Figure S1** | *mTORC1 chains, RMSD convergence plots in MDFF simulations.* | **4** |
| **Figure S2-A, B** | *Evolution of COM distances between protein chains in all four ±RHEB ±ATP mTORC1 complexes at each stage of modelling and simulations* | **5-8** |
| **Figure S3-A, B** | *N-/M-HEAT COM distance comparison between all permutations of RHEB bound and RHEB unbound monomeric units for systems with, without ATP+2$Mg^{2+}$.* | **9-10** |
| **Figure S4-A, B** | *N-/C-Lobe COM distance comparison between kinase domains in all four ±RHEB ±ATP mTORC1 complexes* | **11-12** |
| **Figure S5** | *Model validation with reference to Violin plots depicting distribution of best CASP scores* | **13** |
| **Figure S6** | *Asymmetry convergence analysis and results* | **14-15** |
| **Figure S7** | *Distributions of the change in MMGBSA enthalpy of ATP binding induced by presence of RHEB ΔΔH(±ATP±RHEB) as a function of $r_{cutoff}$ along with decomposition of various contributions* | **16** |
| **Figure S8** | *SASA Time traces(left) and probability distributions (right) across full length trajectories* | **18** |
| **Figure S9** | *Decomposition of ATP-MG contributions to active site conformational heterogeneity.* | **19** |
| **Figure S10** | *Atomic contributions for the first 5 PC modes of SASA based active site conformations* | **20** |
| **Figure S11** | *$\sigma^2_{CVCF}$ traces for individual trajectories of mTORC1 for data shown in Fig 6* | **21** |
| **Figure S12** | *Mean and std.dev $\sigma^2_{CVCF}$ of of mTORC1 as shown in Fig 6 but for -ATP state* | **22** |
| **Figure S13** | *$\sigma^2_{CVCF}$ plots for individual trajectories of mTORC1 for data shown in Fig S12* | **23** |
| **Figure S14** | *Mean and std.dev $\sigma^2_{CVCF}$ of of mTORC1 as shown in Fig 6 for +ATP state (25 ns trajectories)* | **24** |
| **Figure S15** | *$\sigma^2_{CVCF}$ plots for individual trajectories of mTORC1 for data shown in Fig S14* | **25** |
| **Figure S16** | *$\sigma^2_{CVCF}$ traces averages and std-dev over sets of 10×15ns trajectories for the individual protein chains within the four ±RHEB ±ATP mTORC1 complexes.* | **26** |
| **Figure S17** | *$\sigma^2_{CVCF}$ traces (averages, std-dev, and individual trajectory traces) over sets of 10×25ns trajectories for the individual protein chains within the two ±RHEB +ATP mTORC1 complexes.* | **27** |

| Models | Box-Dim. in Å (X*Y*Z) | #Total atoms | #Protein atoms | #Solvent Atoms | #Ions (Na+) | #replicas*length(ns) |
|---|---|---|---|---|---|---|
| mTORC1 | (318.04*245.86*190.21) | 1434169 | 133122 | 1300983 | 64 | 5*15 |
| mTORC1-RHEB | (299.81*242.55*168.10) | 1171082 | 138898 | 1032006 | 74 | 5*15 |
| mTORC1-ATP-Mg | (318.04*245.86*190.21) | 1434205 | 133122 | 1300929 | 64 | 5*15 |
| mTORC1-RHEB-ATP-Mg | (299.81*242.55*168.10) | 1171124 | 138898 | 1031958 | 74 | 5*15 |

**Table S1:** *Details of Solvated models used in Molecular Dynamics Simulations.*

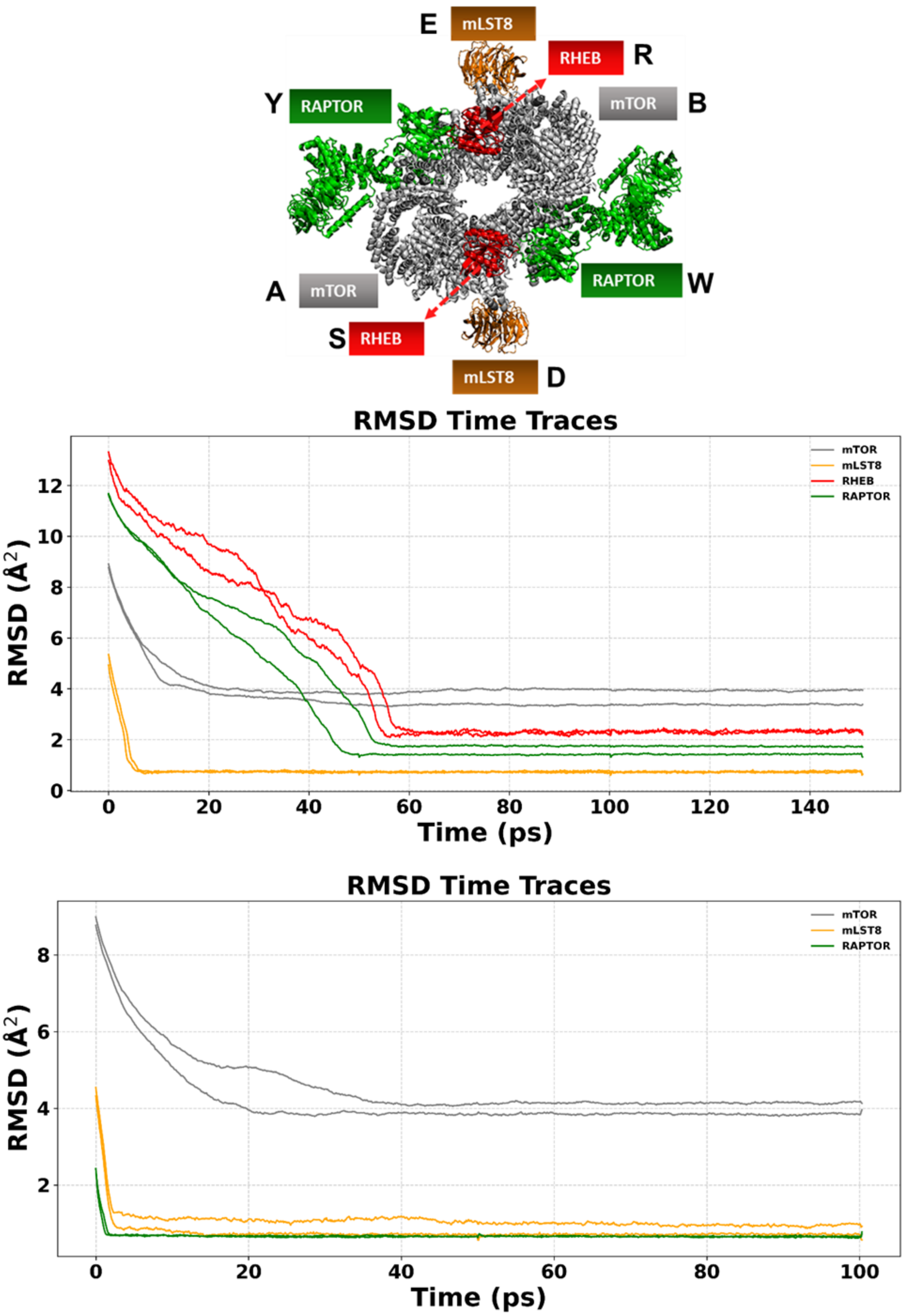


**Figure S1:** *Structure of mTORC1 bound to RHEB annotated with chain ids (Top), RMSD convergence plots for RHEB bound (middle) and RHEB unbound (bottom) due to MDFF simulations.*

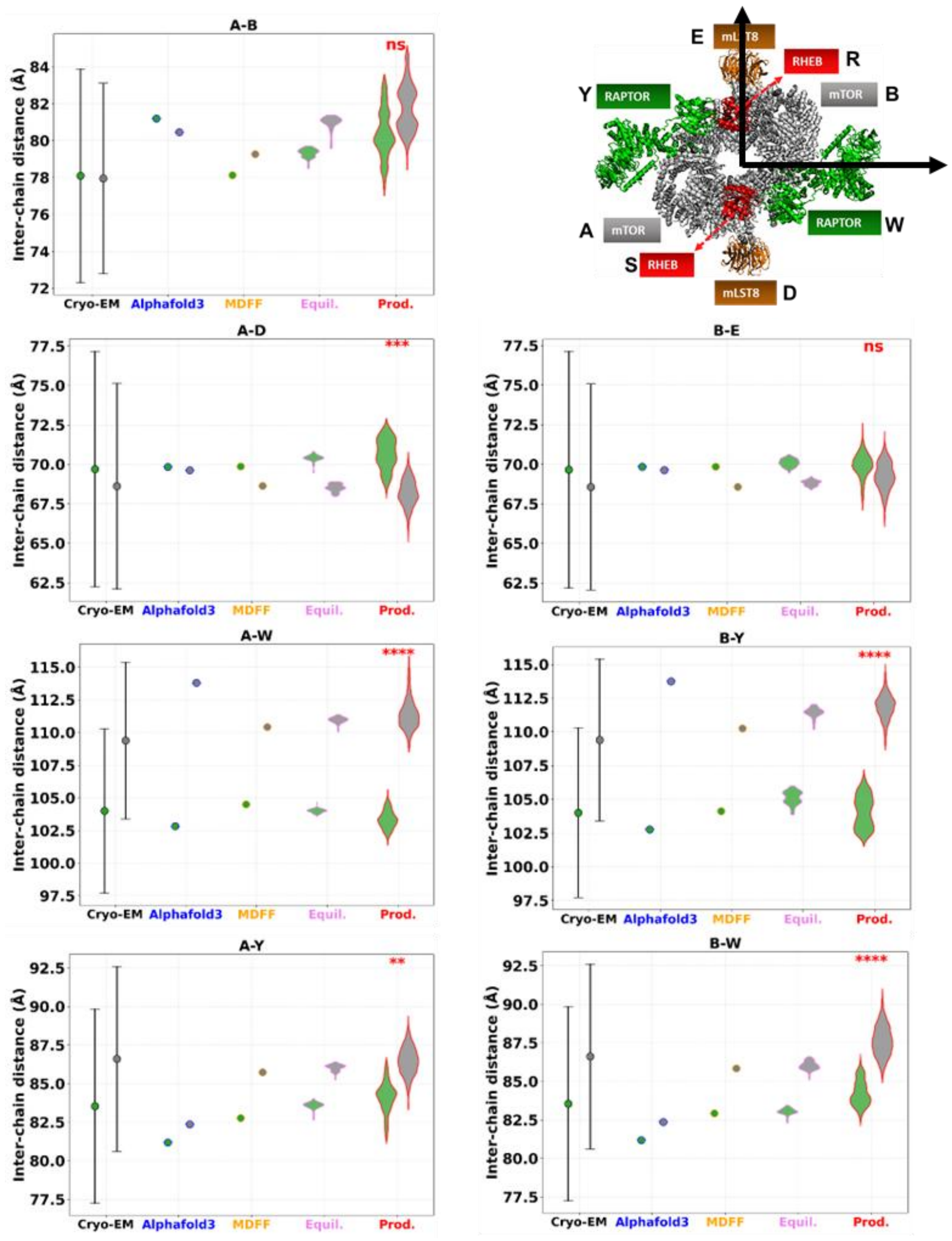


**Figure S2-A:** *COM distances between protein chains for RHEB bound and RHEB unbound mTORC1((with ATP+2Mg*$^{2}$*) at each stage of modelling and simulations compared with Cryo-EM structures*

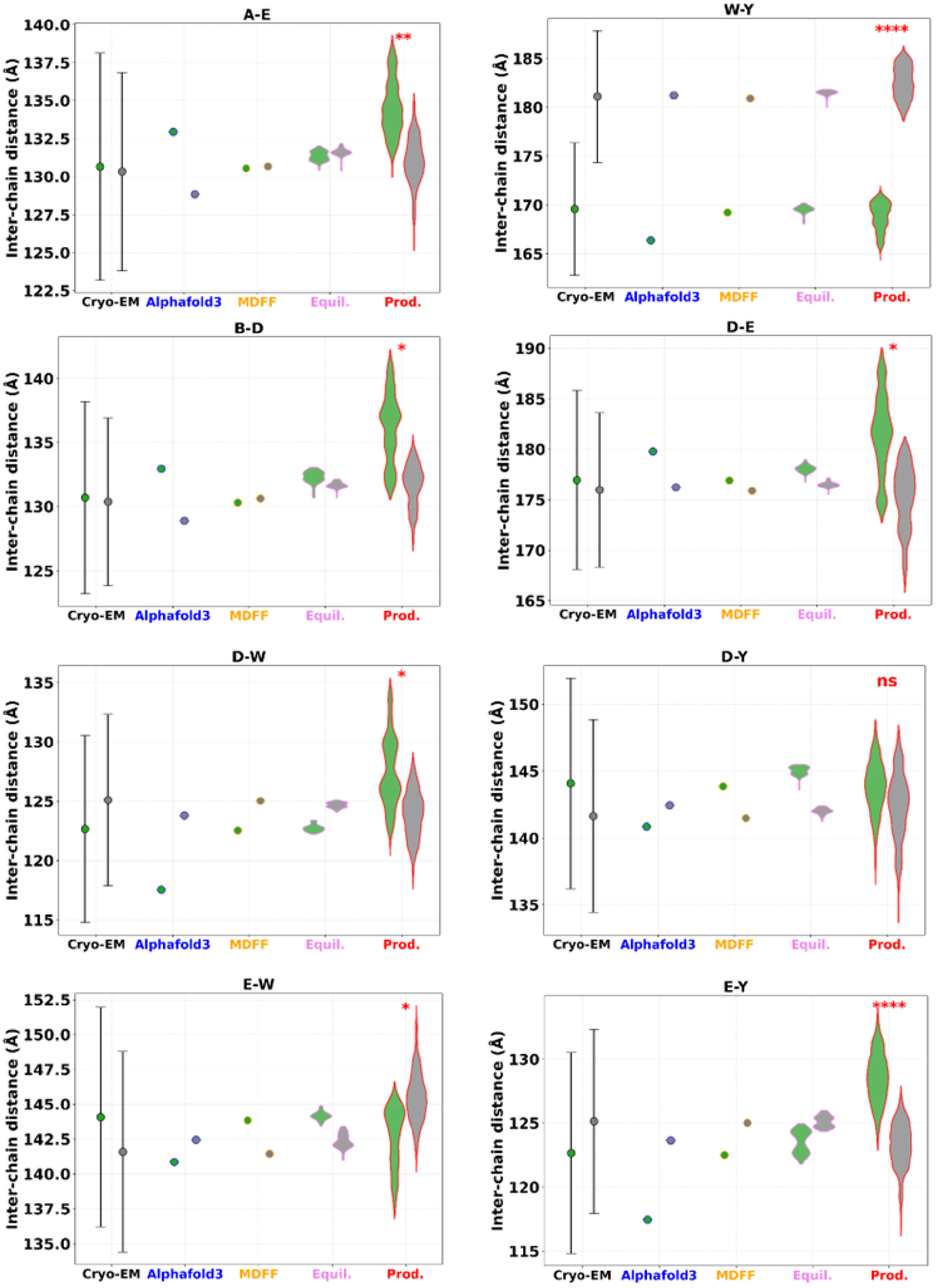


**Figure S2-A (contd.):** *COM distances between protein chains for RHEB bound and RHEB unbound mTORC1((with ATP+2$Mg^{2}$) at each stage of modelling and simulations compared with Cryo-EM structures*

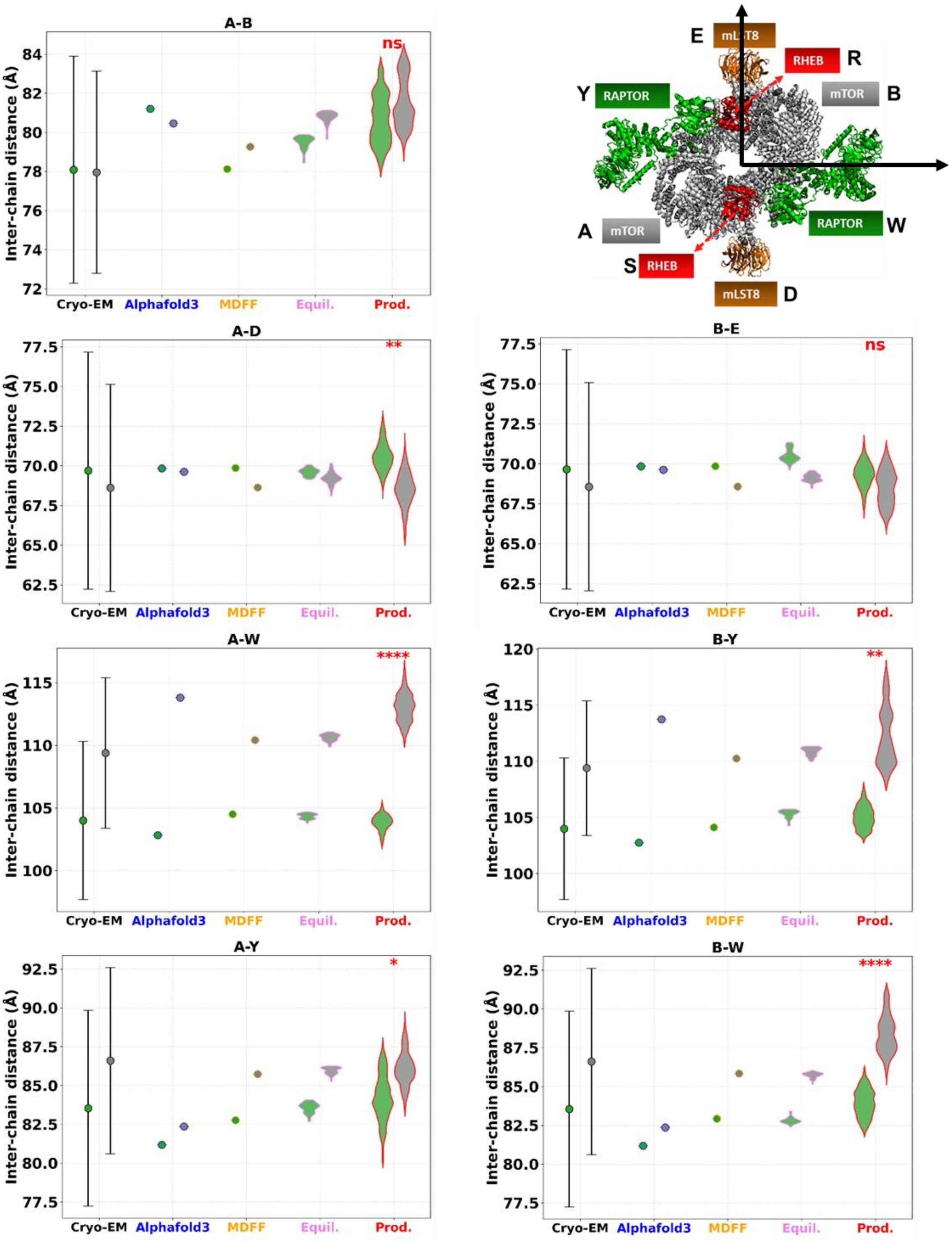


**Figure S2-B:** *COM distances between protein chains for RHEB bound and RHEB unbound mTORC1(without ATP+2Mg2+) at each stage of modelling and simulations compared with Cryo-EM structures*

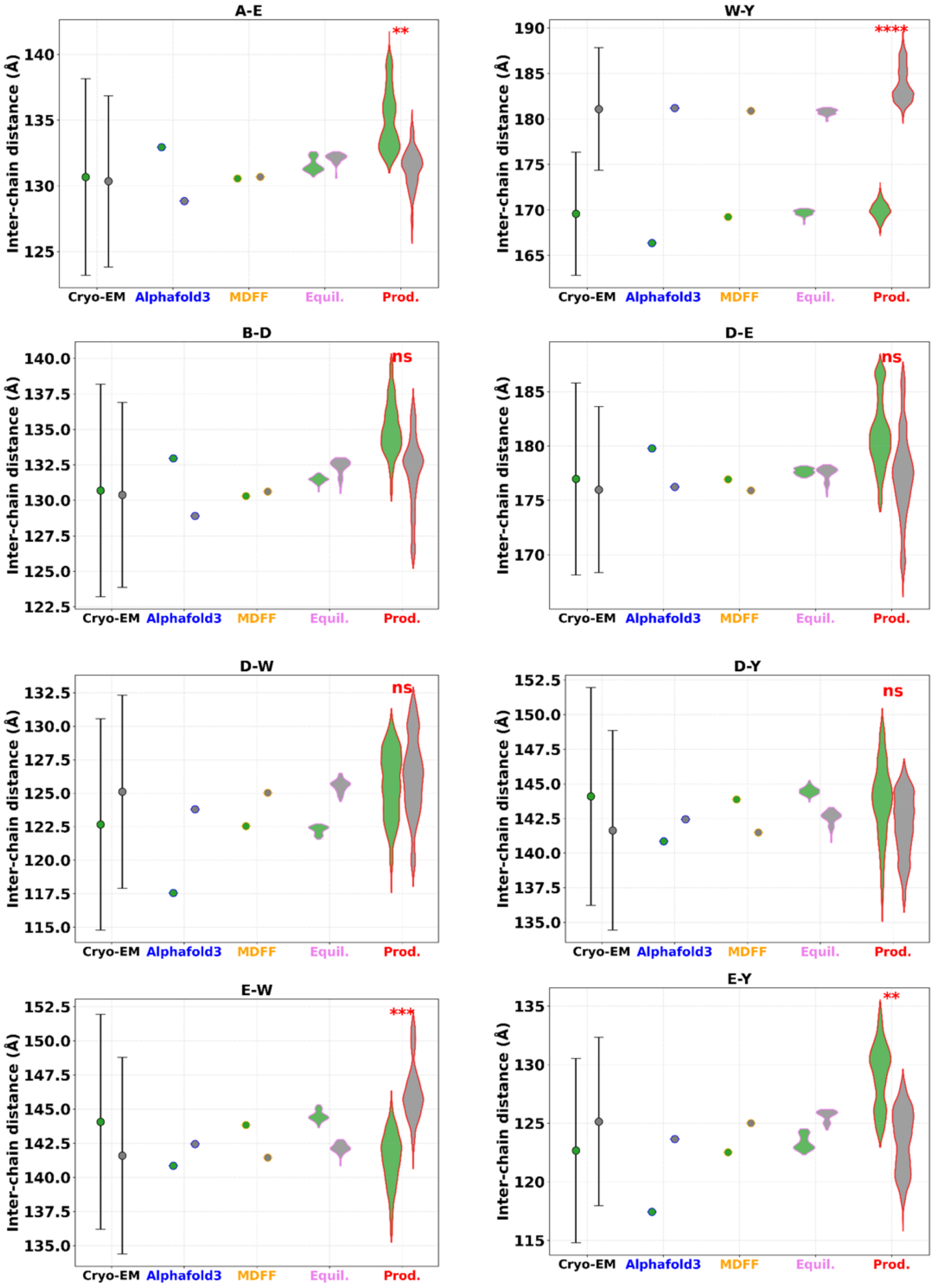


**Figure S2-B (contd.):** *COM distances between protein chains for RHEB bound and RHEB unbound mTORC1(without ATP+2Mg$^{2+}$) at each stage of modelling and simulations compared with Cryo-EM structures*

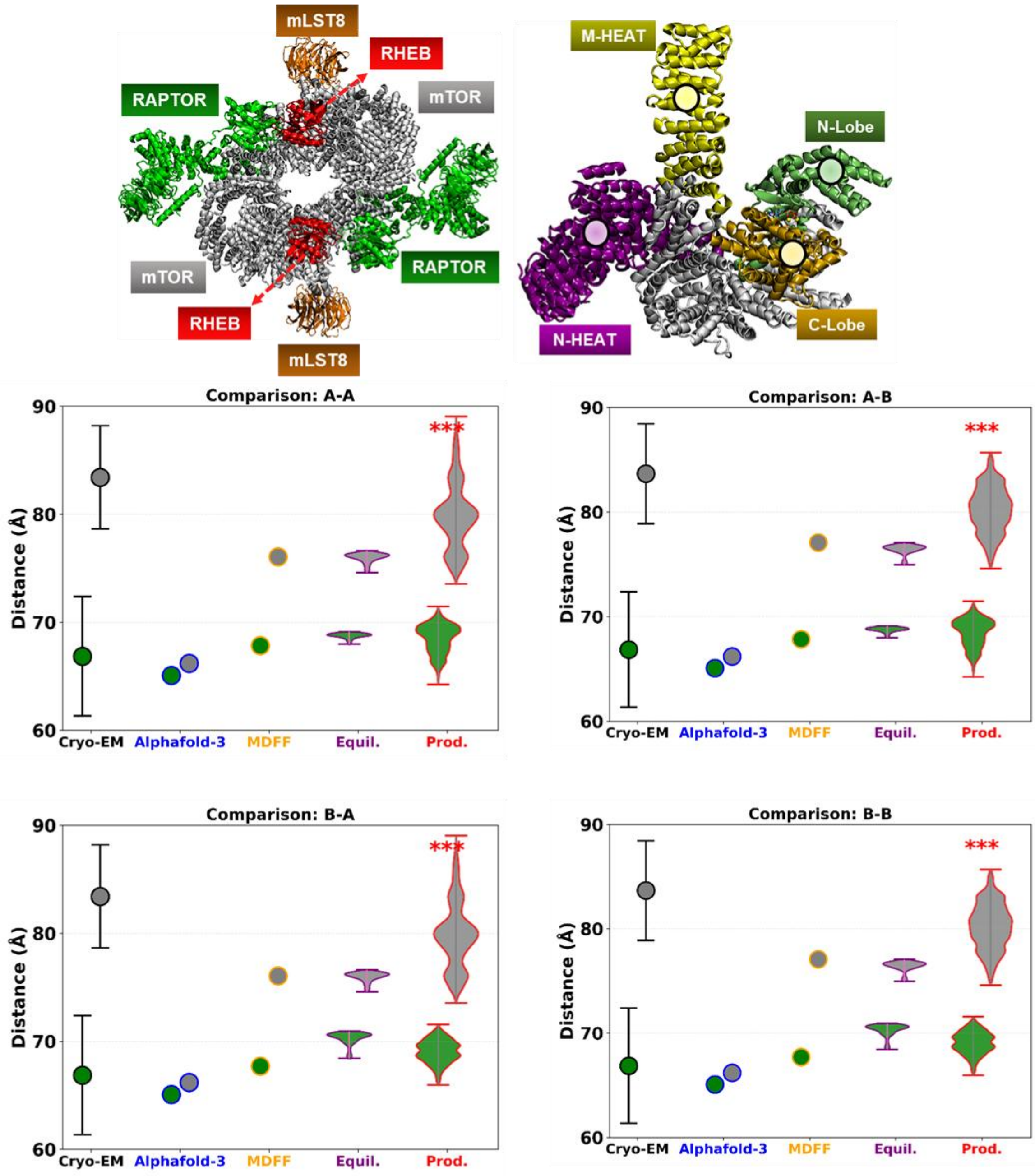


**Figure S3-A:** *N-/M-HEAT COM distance comparison between all permutations of RHEB bound and RHEB unbound monomeric units with ATP+2Mg$^{2+}$.*

| | chainA | chainB |
|---|---|---|
| RHEB bound | 68.77±0.24 | 70.30±0.61 |
| RHEB unbound | 76.00±0.44 | 76.44±0.47 |

**Table S2-A:** *N-/M-HEAT COM distance during Equilibration with ATP+2Mg$^{2+}$*

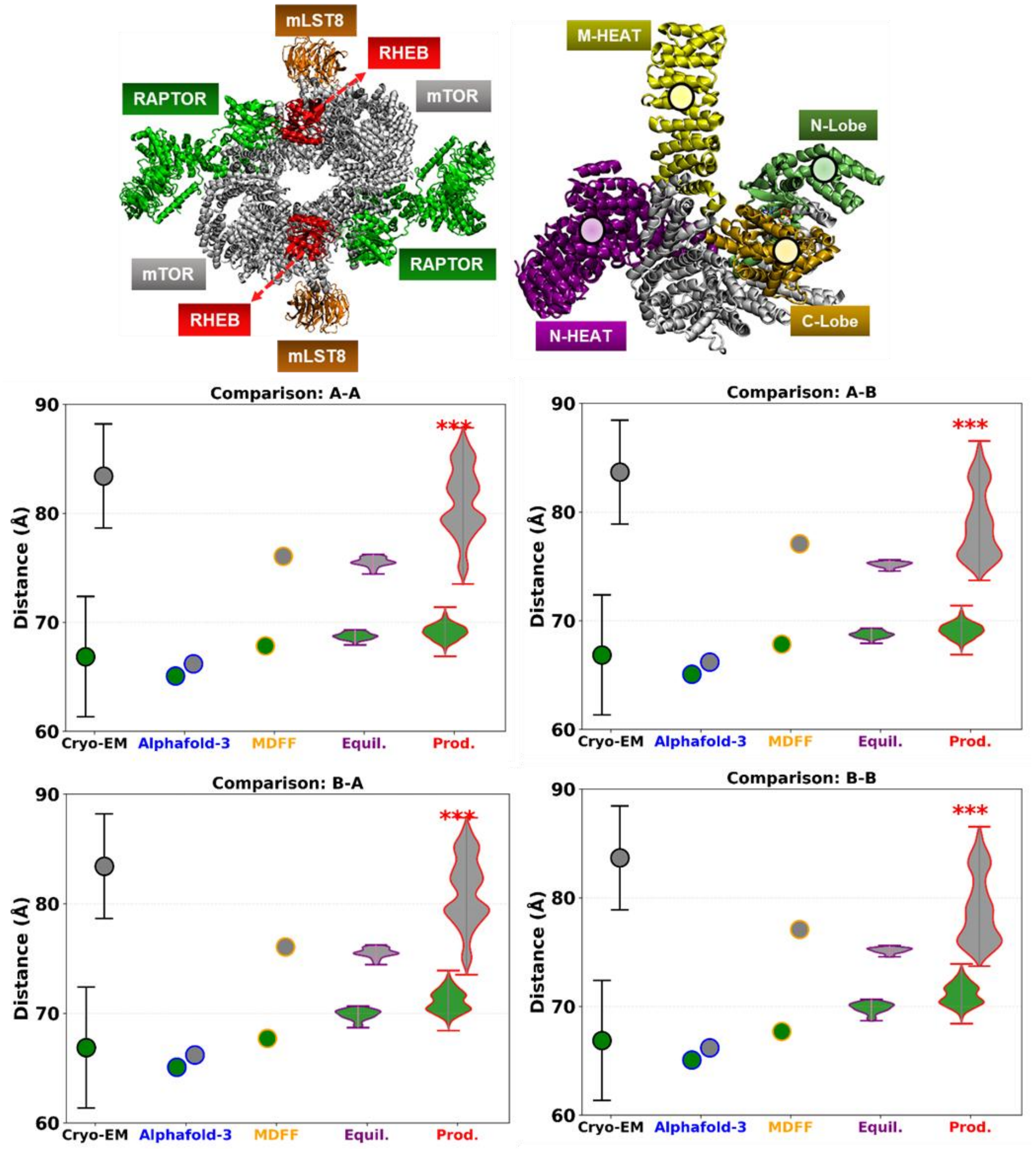


**Figure S3-B:** *N-/M-HEAT COM distance comparison between all permutations of RHEB bound and RHEB unbound monomeric units without ATP+2Mg$^{2+}$.*

| | chainA | chainB |
|---|---|---|
| **RHEB bound** | **68.75±0.30** | **69.90±0.45** |
| **RHEB unbound** | **75.54±0.41** | **75.15±0.24** |

**Table S2-B:** *N-/M-HEAT COM distance during Equilibration without ATP+2Mg$^{2+}$*

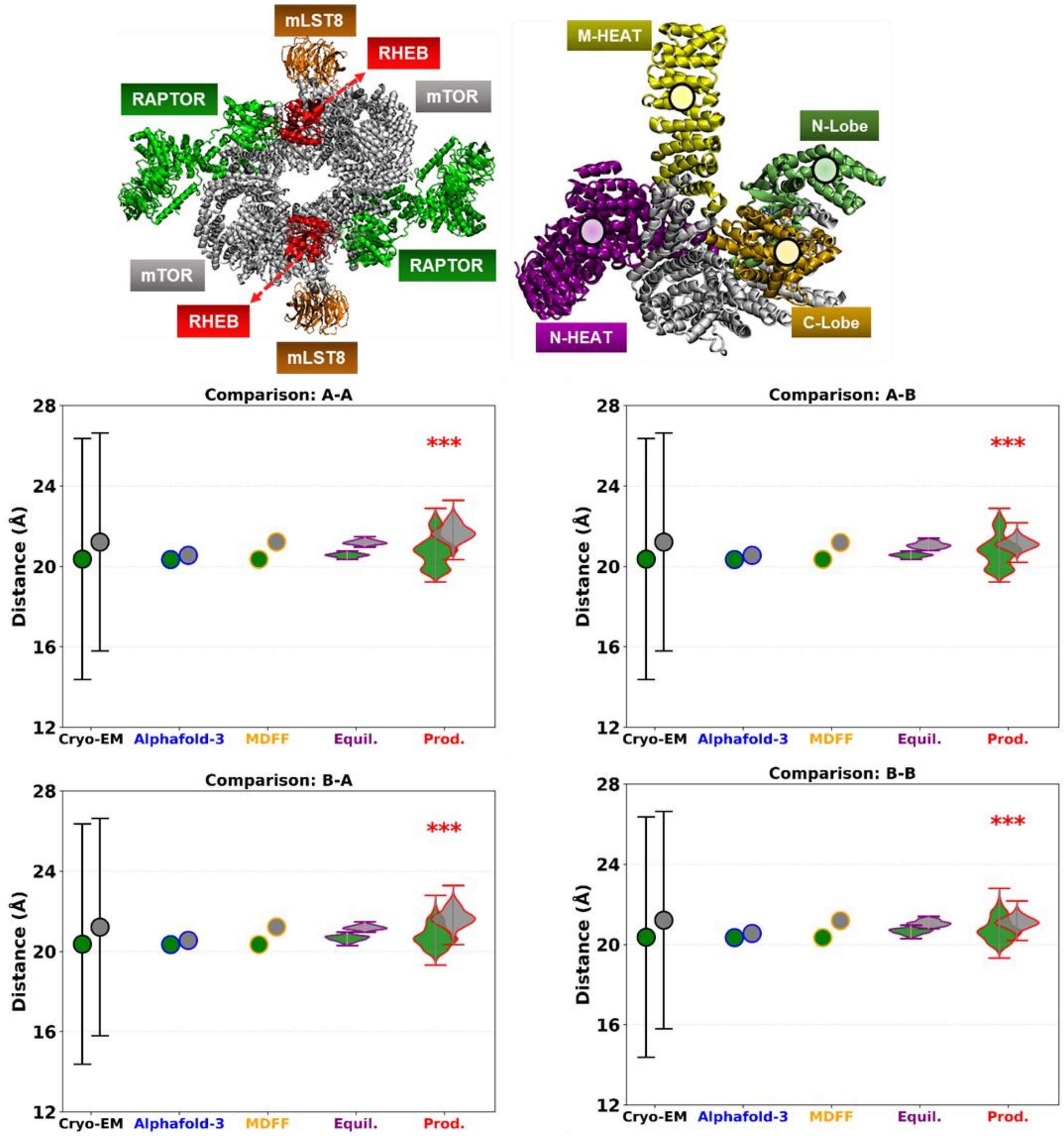


**Figure S4-A:** *N-/C-Lobe COM distance comparison between all permutations of RHEB bound and RHEB unbound monomeric units with ATP+2Mg*$^{2+}$

| | chainA | chainB |
|---|---|---|
| **RHEB-Bound** | **20.55±0.10** | **20.65±0.13** |
| **RHEB-Free** | **21.19±0.10** | **21.06±0.14** |

**Table S3-A:** *N-/C-Lobe COM distance during Equilibration with ATP+2Mg*$^{2+}$

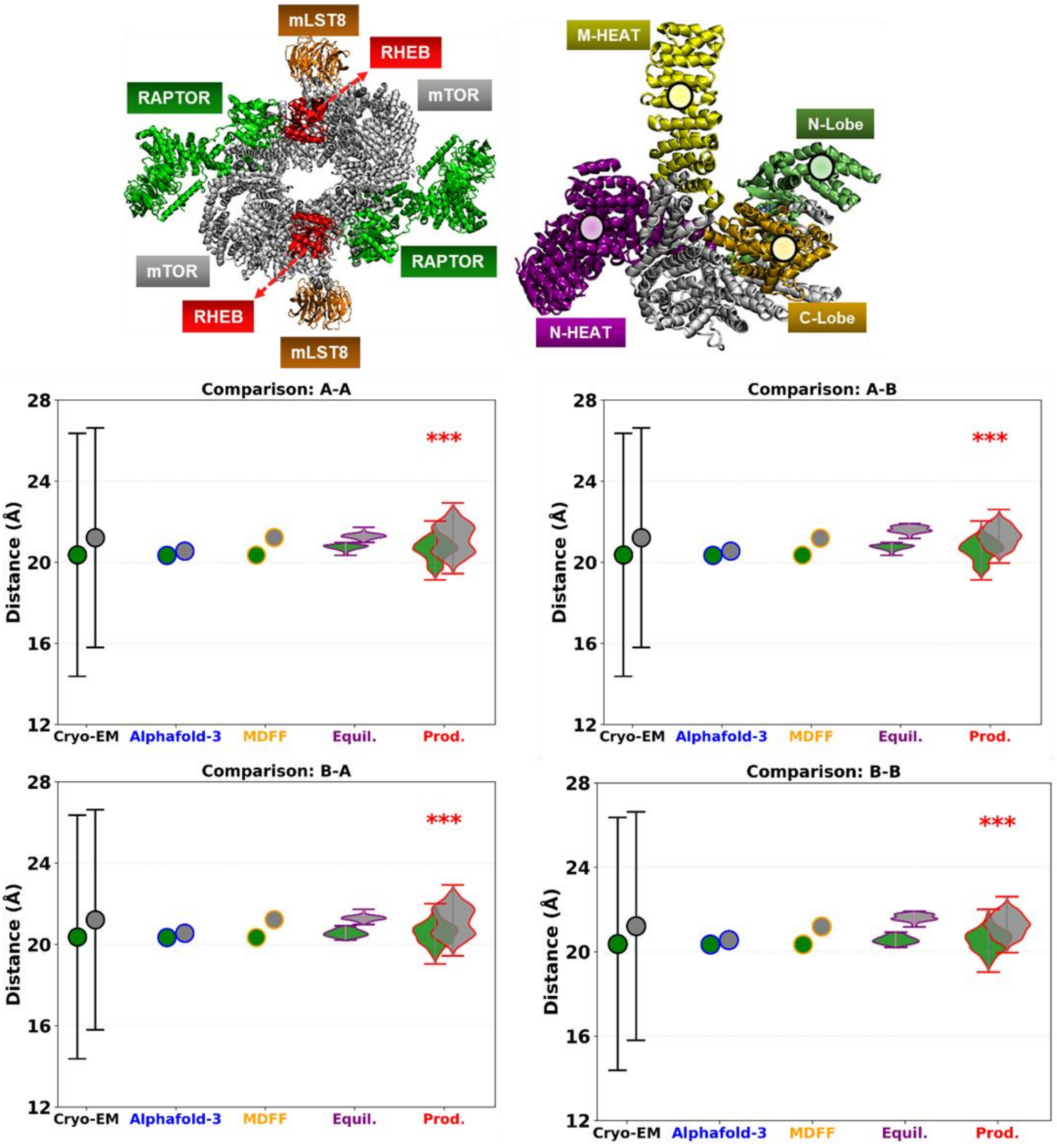


**Figure S4-B:** *N-/C-Lobe COM distance comparison between all permutations of RHEB bound and RHEB unbound monomeric units without ATP+2Mg*$^{2+}$

| | chainA | chainB |
|---|---|---|
| RHEB-Bound | 20.74±0.12 | 20.56±0.17 |
| RHEB-Free | 21.29±0.12 | 21.63±0.15 |

**Table S3-B:** *N-/C-Lobe COM distance during Equilibration without ATP+2Mg*$^{2}$

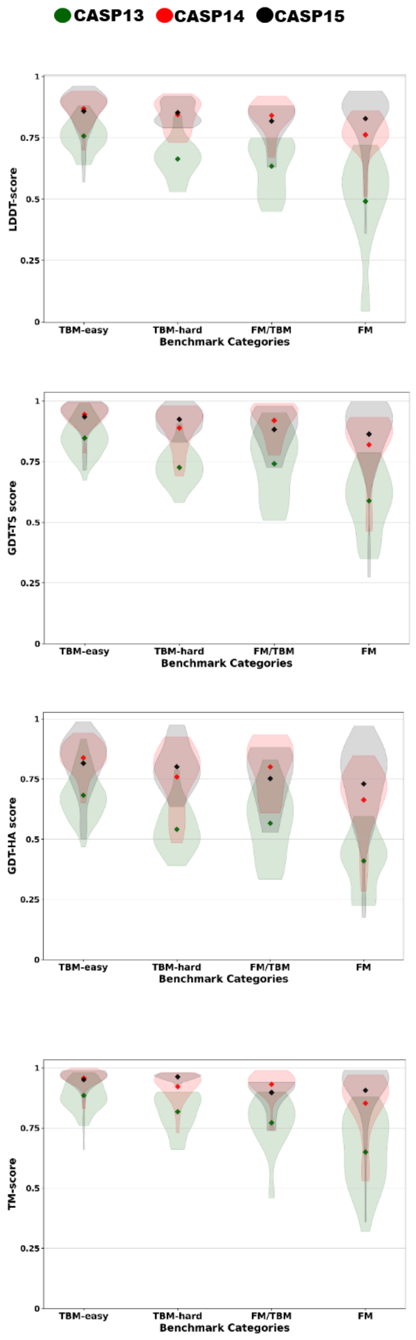


**RHEB-Bound**

| Chain | AF3 | MDFF | Equ |
|---|---|---|---|
| A | 0.94 | 0.87 | 0.84±0.01 |
| B | 0.94 | 0.91 | 0.84±0.00 |
| D | 0.91 | 0.93 | 0.87±0.01 |
| E | 0.91 | 0.93 | 0.86±0.01 |
| W | 0.92 | 0.93 | 0.86±0.00 |
| Y | 0.92 | 0.92 | 0.84±0.01 |
| R | 0.91 | 0.93 | 0.78±0.01 |
| S | 0.91 | 0.94 | 0.76±0.01 |

**RHEB-Free**

| Chain | AF3 | MDFF | Equ |
|---|---|---|---|
| A | 0.95 | 0.89 | 0.85±0.00 |
| B | 0.95 | 0.89 | 0.85±0.01 |
| D | 0.92 | 0.94 | 0.87±0.01 |
| E | 0.92 | 0.93 | 0.86±0.01 |
| W | 0.94 | 0.93 | 0.88±0.01 |
| Y | 0.94 | 0.93 | 0.89±0.01 |

| Chain | AF3 | MDFF | Equ |
|---|---|---|---|
| A | 0.88 | 0.86 | 0.71±0.03 |
| B | 0.88 | 0.92 | 0.74±0.03 |
| D | 0.96 | 0.98 | 0.92±0.01 |
| E | 0.96 | 0.98 | 0.91±0.01 |
| W | 0.88 | 0.97 | 0.86±0.02 |
| Y | 0.88 | 0.97 | 0.81±0.03 |
| R | 0.96 | 0.97 | 0.83±0.01 |
| S | 0.96 | 0.97 | 0.76±0.02 |

| Chain | AF3 | MDFF | Equ |
|---|---|---|---|
| A | 0.53 | 0.87 | 0.49±0.02 |
| B | 0.53 | 0.88 | 0.48±0.01 |
| D | 0.96 | 0.99 | 0.91±0.01 |
| E | 0.96 | 0.98 | 0.89±0.02 |
| W | 0.88 | 0.98 | 0.89±0.01 |
| Y | 0.88 | 0.98 | 0.87±0.02 |

| Chain | AF3 | MDFF | Equ |
|---|---|---|---|
| A | 0.69 | 0.76 | 0.48±0.04 |
| B | 0.69 | 0.85 | 0.51±0.04 |
| D | 0.83 | 0.91 | 0.74±0.02 |
| E | 0.83 | 0.91 | 0.72±0.02 |
| W | 0.72 | 0.90 | 0.66±0.03 |
| Y | 0.72 | 0.88 | 0.59±0.03 |
| R | 0.85 | 0.88 | 0.63±0.02 |
| S | 0.85 | 0.91 | 0.55±0.03 |

| Chain | AF3 | MDFF | Equ |
|---|---|---|---|
| A | 0.37 | 0.78 | 0.28±0.02 |
| B | 0.36 | 0.79 | 0.28±0.01 |
| D | 0.87 | 0.94 | 0.73±0.02 |
| E | 0.87 | 0.91 | 0.70±0.02 |
| W | 0.72 | 0.91 | 0.70±0.02 |
| Y | 0.72 | 0.91 | 0.67±0.03 |

| Chain | AF3 | MDFF | Equilibrated |
|---|---|---|---|
| A | 0.99 | 0.96 | 0.96±0.00 |
| B | 0.99 | 0.98 | 0.95±0.00 |
| D | 0.99 | 0.99 | 0.97±0.00 |
| E | 0.99 | 0.99 | 0.97±0.00 |
| W | 0.99 | 0.99 | 0.98±0.00 |
| Y | 0.99 | 0.99 | 0.97±0.00 |
| R | 0.97 | 0.97 | 0.88±0.01 |
| S | 0.97 | 0.98 | 0.86±0.01 |
| FC | 0.88 | 0.99 | 0.99±0.00 |

| Chain | AF3 | MDFF | Equ |
|---|---|---|---|
| A | 0.85 | 0.96 | 0.88±0.0 |
| B | 0.85 | 0.96 | 0.87±0.0 |
| D | 0.98 | 0.99 | 0.97±0.0 |
| E | 0.98 | 0.99 | 0.96±0.0 |
| W | 0.99 | 0.99 | 0.99±0.0 |
| Y | 0.99 | 0.99 | 0.99±0.0 |
| FC | 0.93 | 0.99 | 0.98±0.0 |

**Figure S5: (Left)** *Violins depicting distribution of best CASP scores of each category for model validation metrics.*

**Table S4: (Right)** *Model validation metrics scores with errors.*

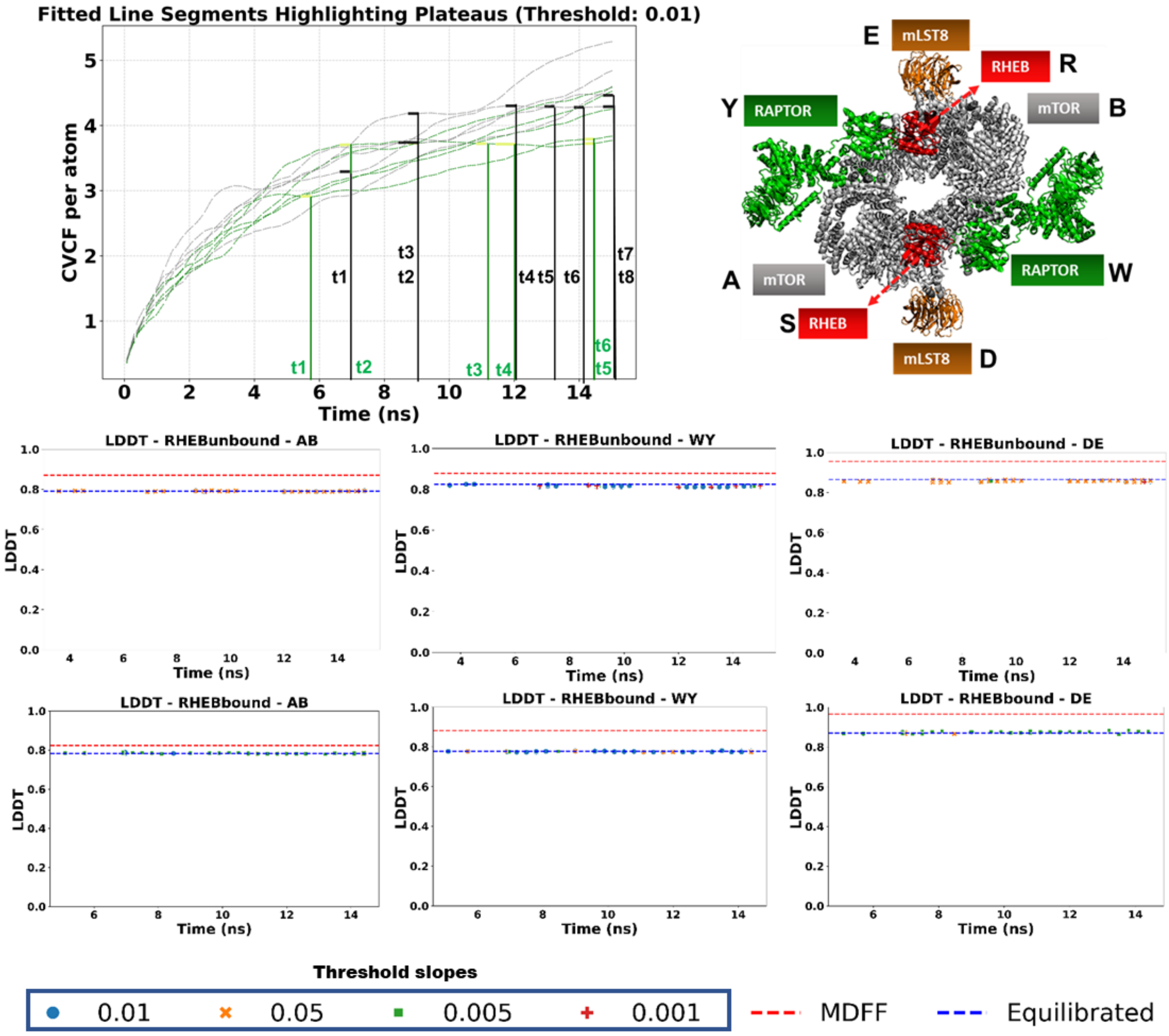


**Figure S6:** *Asymmetry convergence analysis and results*

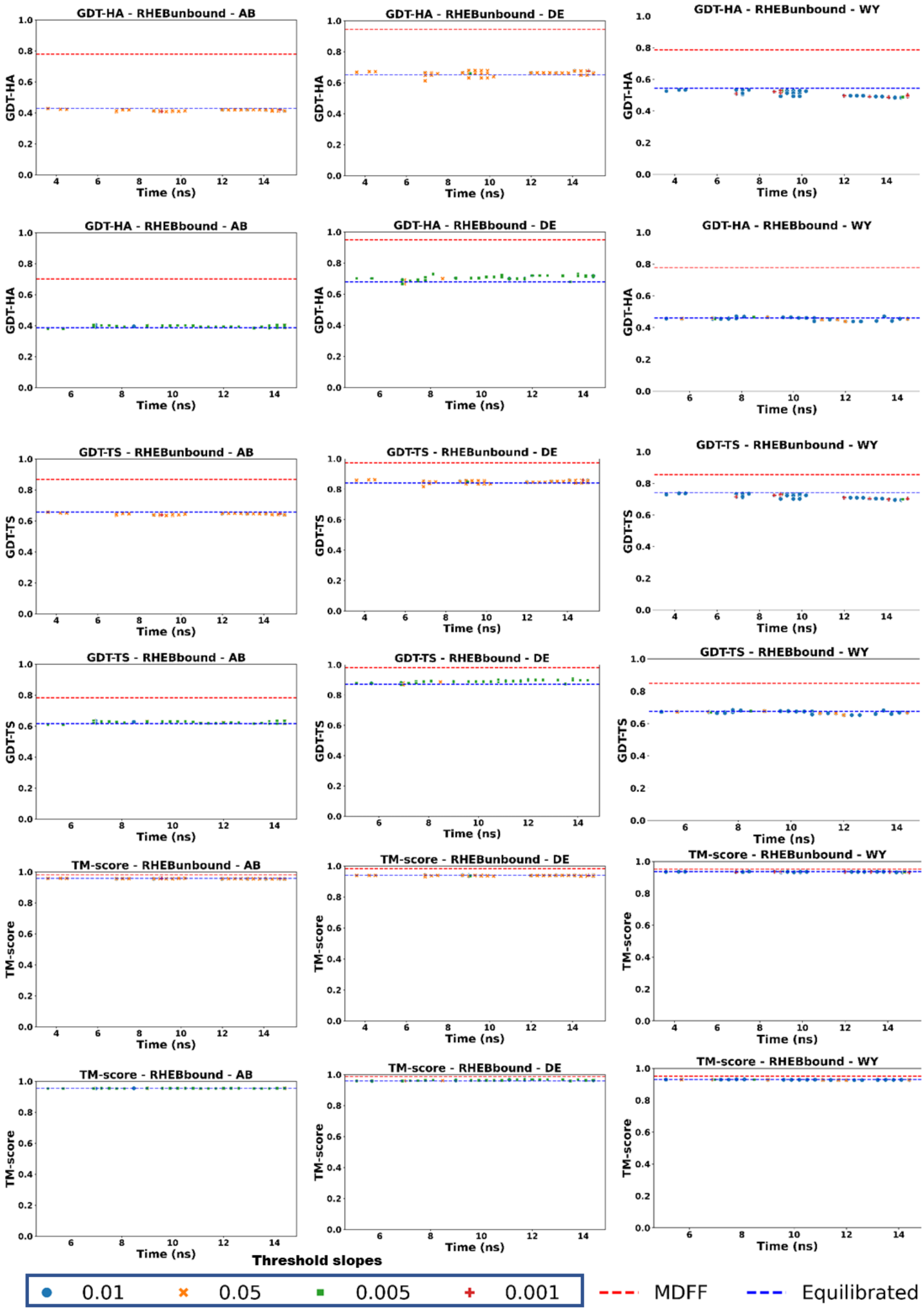


**Figure S6(contd.):** *Asymmetry convergence analysis and results*

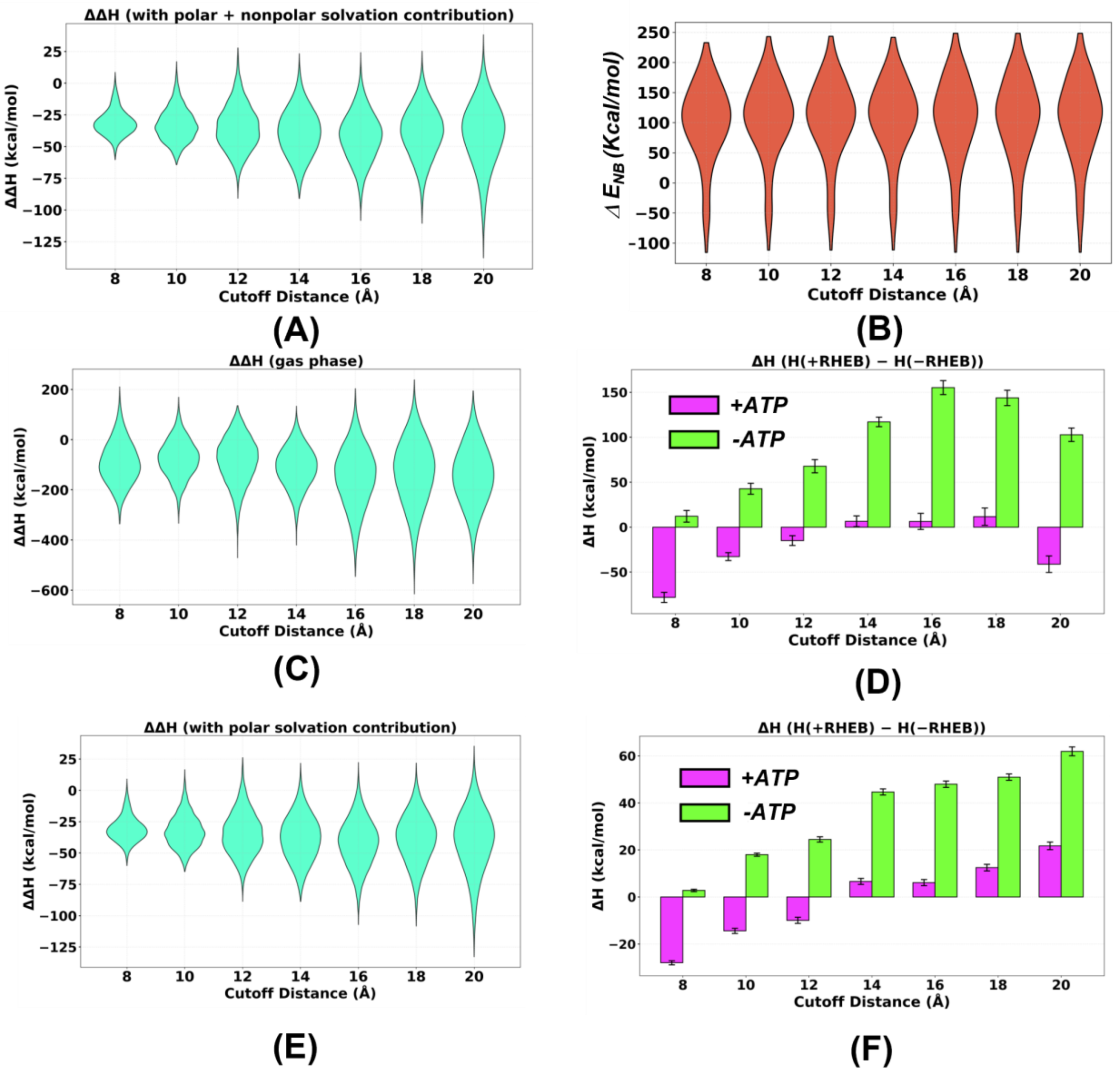


**Figure S7:** (A/C/E) *Distributions of the change in enthalpy of ATP binding (**Eqn.11**) induced by presence of RHEB ΔΔH(±ATP±RHEB) as a function of* $r_{cutoff}$*. Distributions are shown for the MMGBSA enthalpy (**Eqn.11**) with (A) All terms, (C) only gas phase terms, and (E) excluding the non-polar solvation. (B) Distributions of the change in non-bonded interaction energy (*$\Delta E_{NB}$ *) between ATP+2*$Mg^{2+}$ *and surrounding residues induced by RHEB binding. (D/F) Change in the enthalpy of mTORC1 (ΔH(±RHEB)) induced by RHEB binding in the presence and absence of ATP. Data is shown for MMGBSA enthalpy (**Eqn.11**) with (D) only gas phase terms, and (F) excluding the non-polar solvation.*

| Energy terms | 8A | 10A | 12A | 14A | 16A | 18A | 20A |
|---|---|---|---|---|---|---|---|
| BOND | 26.91± 0.99 | 26.98± 1.08 | 26.83± 1.29 | 26.78± 1.24 | 27.00± 1.01 | 26.98± 1.04 | 27.10± 1.31 |
| ANGLE | 57.64± 2.55 | 56.90± 3.91 | 56.33± 4.35 | 55.37± 5.49 | 54.79 ± 5.67 | 55.53± 6.78 | 56.94± 7.47 |
| DIHED | 37.47± 3.96 | 37.22± 4.58 | 38.60± 5.69 | 37.78± 7.54 | 37.45± 10.06 | 36.83± 10.76 | 38.80± 12.19 |
| IMPRP | 0.60± 0.34 | 0.64± 0.45 | 0.60± 0.52 | 0.60± 0.53 | 0.56± 0.52 | 0.61± 0.68 | 0.61± 0.64 |
| ELECT | -2028.68± 71.49 | -1991.90± 59.34 | -1985.59± 62.04 | -2016.89± 62.24 | -2035.16± 101.46 | -2010.99± 99.43 | -2015.99± 90.41 |
| VDW | 42.43± 5.12 | 42.25± 4.47 | 44.12± 5.27 | 43.21± 6.46 | 43.47± 9.78 | 42.89± 11.90 | 46.48± 13.11 |
| POTENTIAL | -1863.61± 70.84 | -1827.91± 60.17 | -1819.12± 62.09 | -1853.15± 59.75 | -1871.89± 96.93 | -1848.15± 95.92 | -1846.05± 89.62 |
| POLAR | 54.99± 69.72 | 18.74± 61.77 | 10.73± 57.46 | 38.66± 58.98 | 53.00± 96.27 | 32.83± 92.47 | 31.65± 83.63 |
| NONPOLAR | -2.85± 0.47 | -2.73± 0.53 | -2.75± 0.78 | -3.16± 0.80 | -3.00± 1.01 | -2.85± 1.18 | -2.62± 1.42 |

| Energy terms | 8A | 10A | 12A | 14A | 16A | 18A | 20A |
|---|---|---|---|---|---|---|---|
| BOND | 25.08± 0.57 | 25.03± 0.66 | 25.12± 0.91 | 25.23± 1.02 | 25.22± 1.11 | 25.26± 1.24 | 25.44± 1.17 |
| ANGLE | 51.32± 4.43 | 52.01± 4.72 | 52.80± 5.05 | 52.58± 5.41 | 50.76± 5.32 | 50.78± 5.62 | 50.47± 6.30 |
| DIHED | 34.75± 4.11 | 33.44± 4.52 | 33.88± 4.91 | 32.27± 5.22 | 31.46± 6.57 | 31.95± 6.25 | 30.92± 5.89 |
| IMPRP | 0.61± 0.28 | 0.61± 0.30 | 0.64± 0.36 | 0.63± 0.41 | 0.71± 0.45 | 0.71± 0.48 | 0.79± 0.52 |
| ELECT | -1918.32± 47.97 | -1894.00± 43.90 | -1882.18± 64.98 | -1884.46± 51.22 | -1859.27± 69.12 | -1852.36± 88.72 | -1834.49± 82.88 |
| VDW | 33.20± 5.66 | 30.47± 7.71 | 33.33± 7.49 | 31.11± 8.75 | 28.04± 12.04 | 27.61± 12.74 | 24.74± 14.08 |
| POTENTIAL | -1773.35± 48.45 | -1752.44± 43.55 | -1736.41± 67.15 | -1742.65± 53.49 | -1723.08± 66.77 | -1716.05± 85.78 | -1702.13± 76.80 |
| POLAR | -4.51± 47.48 | -24.30± 45.02 | -37.58± 63.60 | -33.81± 50.16 | -53.94± 65.32 | -60.79± 83.64 | -72.07± 78.96 |
| NONPOLAR | -2.53± 0.53 | -2.59± 0.67 | -2.36± 0.80 | -2.55± 0.84 | -2.80± 1.12 | -2.67± 1.26 | -2.54± 1.47 |

**Table S5:** *Individual Energy contributions to the Enthalpy change for RHEB bound(green) and RHEB unbound (grey).*

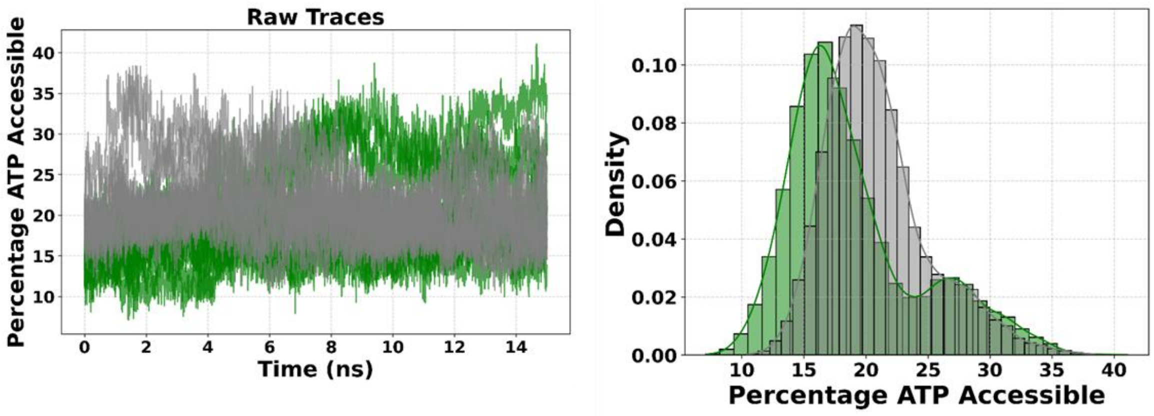


**Figure S8:** *SASA Time traces(left) and probability distribution(right) across full length trajectories*

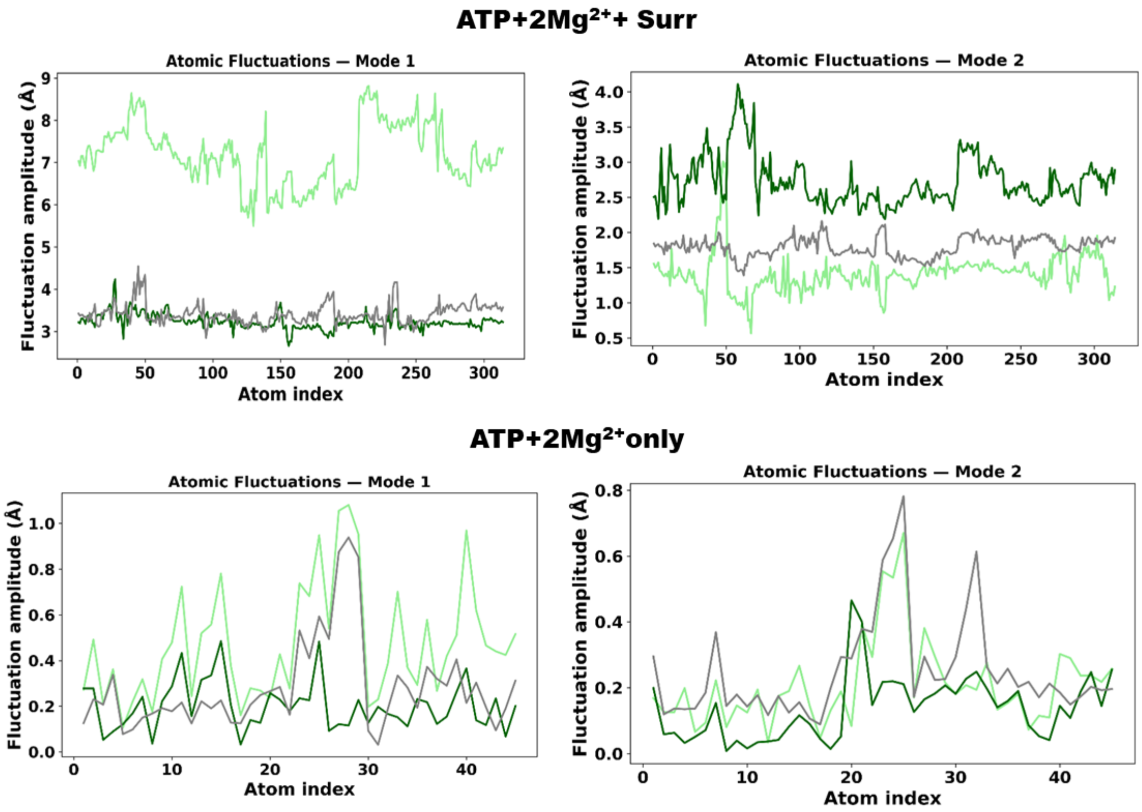


**Figure S9:** *ATP-MG contribution is less to the conformational heterogeneity. We obtain this by performing PCA on ATP+2Mg$^{2+}$+surrounding(top) as well as only ATP+2Mg$^{2+}$+(bottom). We conclude that the major differences between mode1 and mode 2 of the active site arise predominantly due to surrounding residues and not ATP-2Mg$^{2+}$*

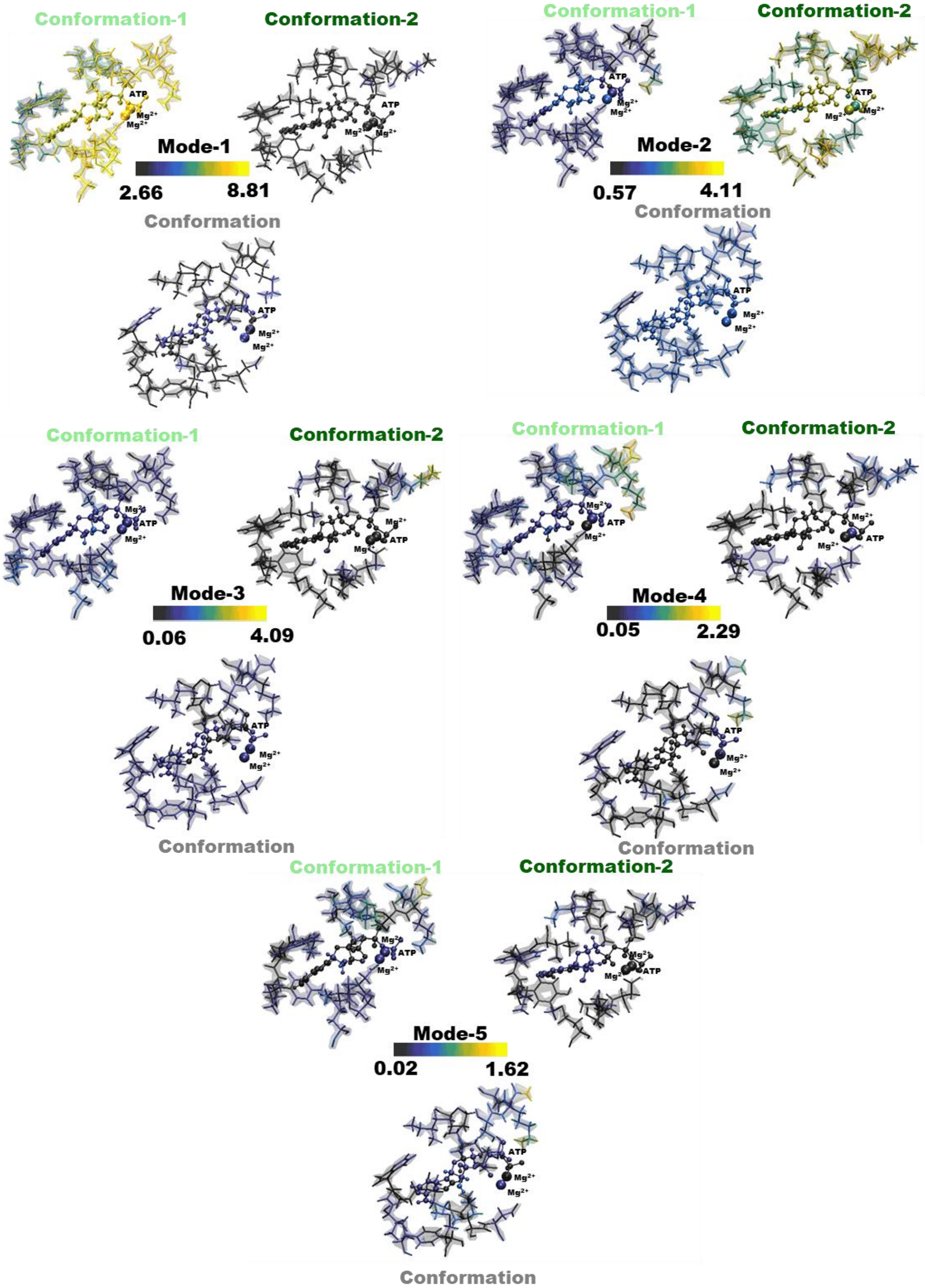


**Figure S10:** *Atomic contributions to first 5 PC modes of SASA based Active site conformations*

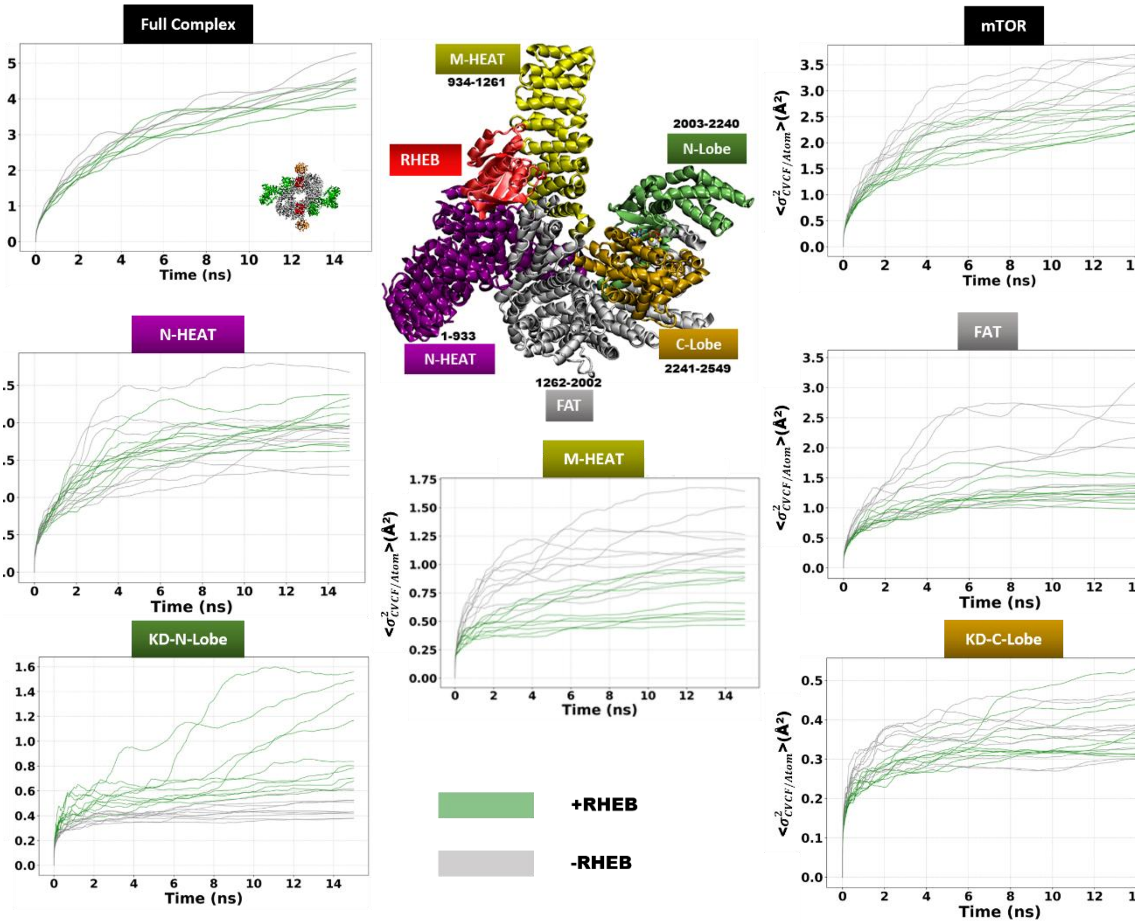


**Figure S11:** *Per atom $\sigma^2{}_{CVCF}$ traces of individual trajectories for the data in Figure 6 of the main manuscript*

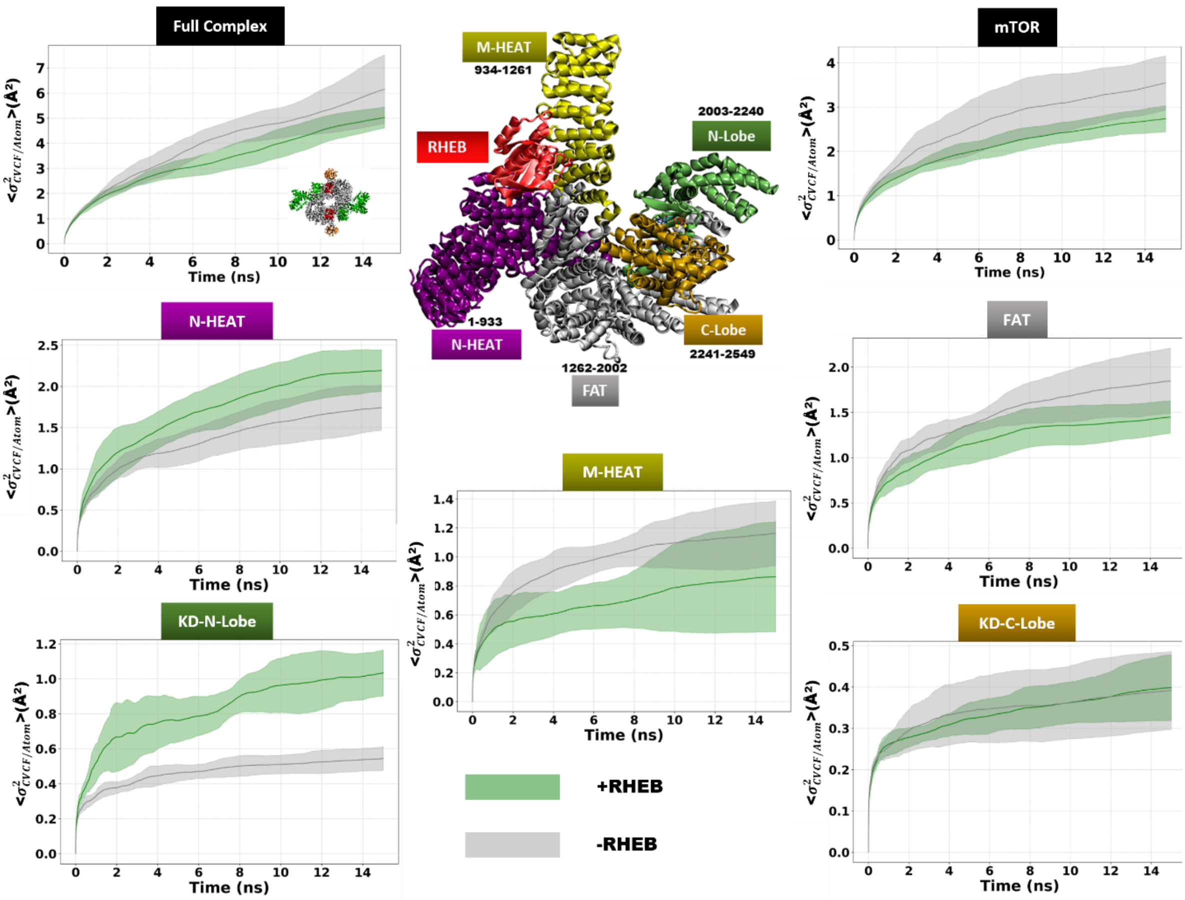


**Figure S12:** *Mean and std.dev of per atom $\sigma^2_{CVCF}$ traces of structured $C_\alpha$ atoms of Full Complex (top left) along with mTOR (top right) along with plots for the same for respective domains constituting mTOR without ATP from 5 × 15ns MD trajectories for full mTORC1 and 10 × 15ns (two instances for each complex) for mTOR chains and respective domains with (green) and without (grey) RHEB. The inset structure in the top row shows the mTOR chain along with its constituent domains bound to RHEB.*

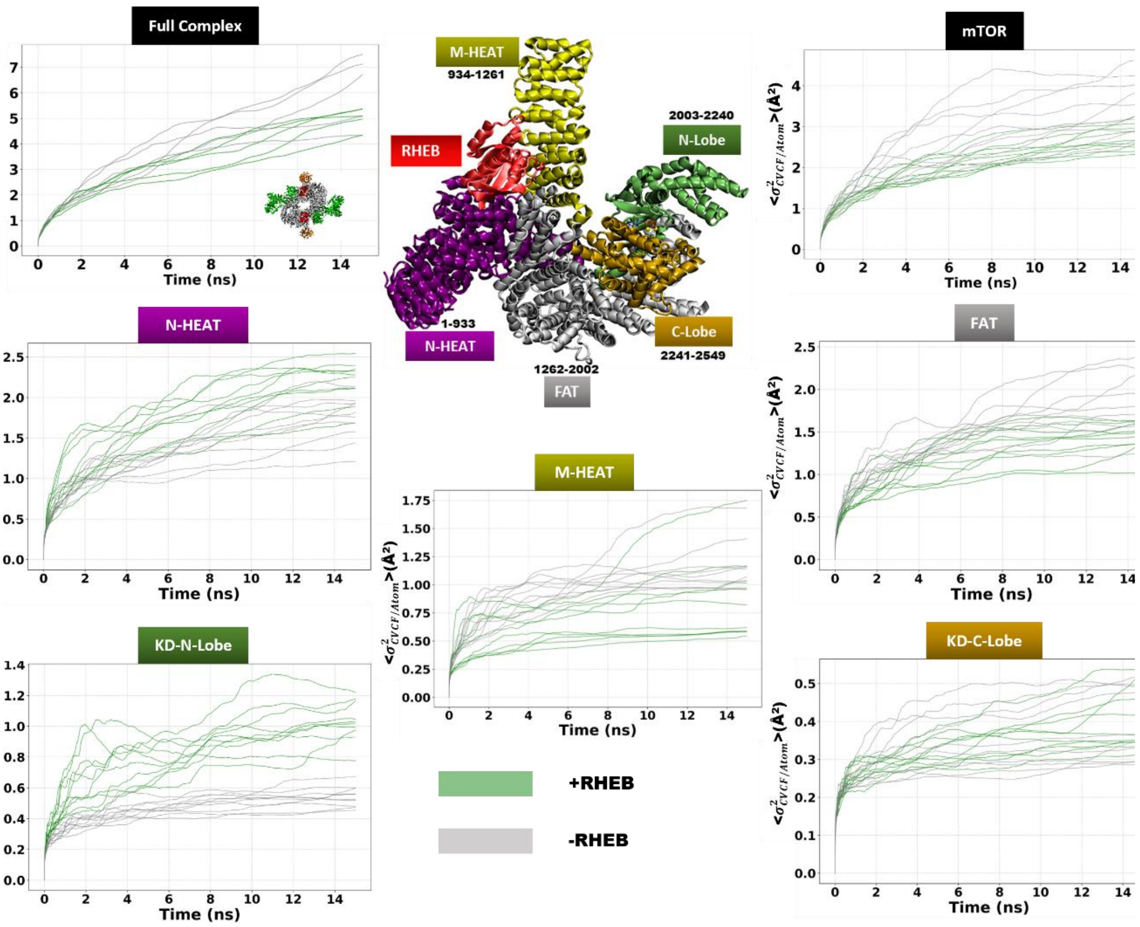


**Figure S13:** *Per atom $\sigma^2_{CVCF}$ traces of individual trajectories for the data in Figure S12*

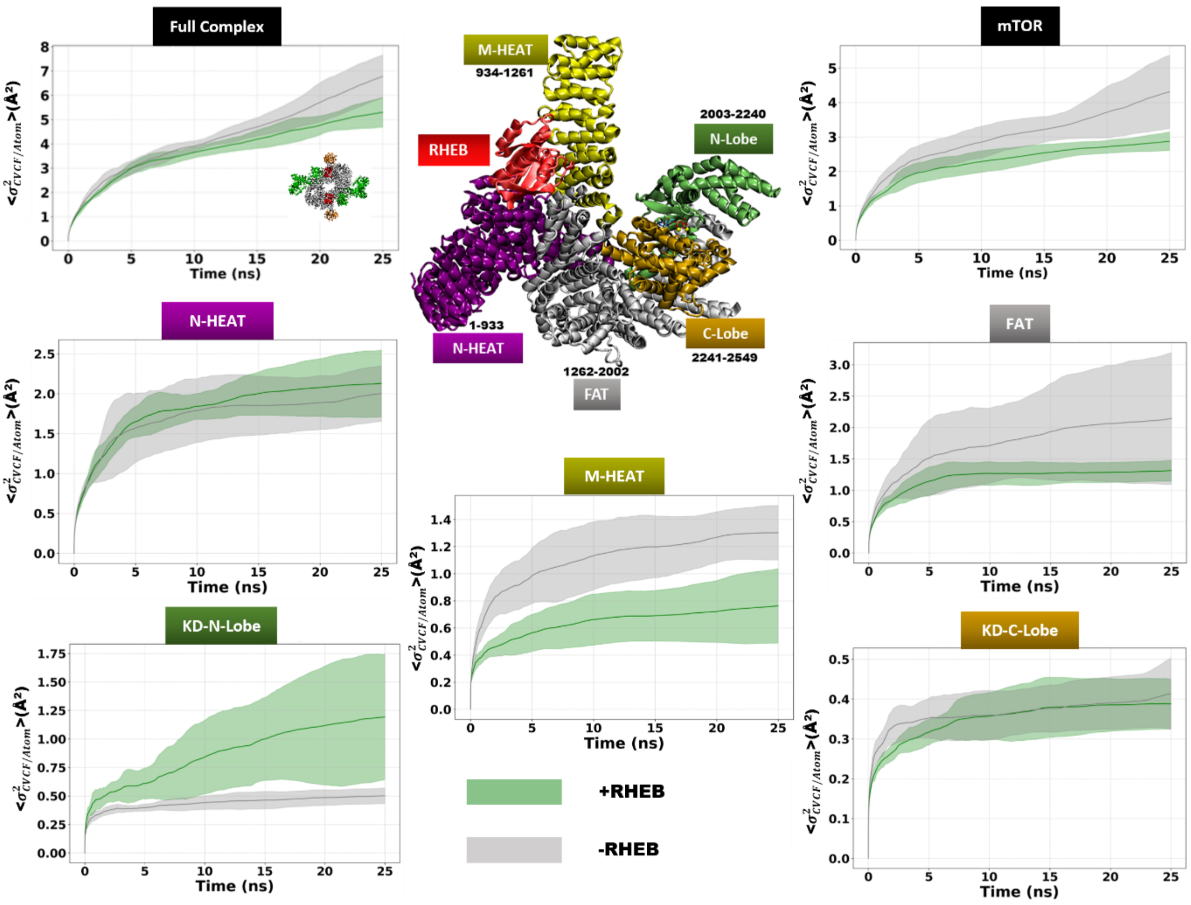


**Figure S14:** *Mean and std.dev of per atom* $\sigma^2_{CVCF}$ *traces of structured* $C_\alpha$ *atoms of Full Complex (top left) along with mTOR (top right) along with plots for the same for respective domains constituting mTOR without ATP from 5 × 25ns MD trajectories for full mTORC1 and 10 × 25ns (two instances for each complex) for mTOR chains and respective domains with (green) and without (grey) RHEB. The inset structure in the top row shows the mTOR chain along with its constituent domains bound to RHEB.*

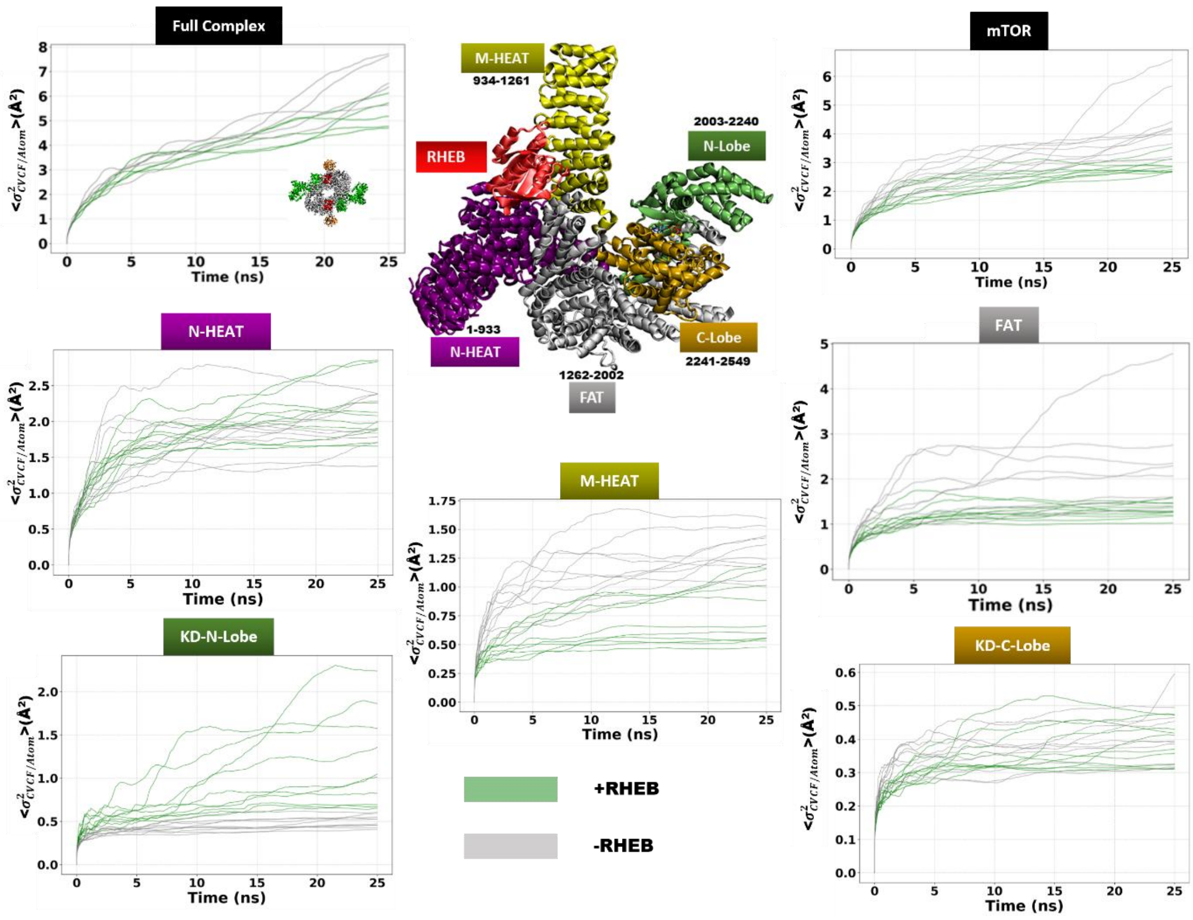


**Figure S15:** *Per atom* $\sigma^2_{CVCF}$ *traces of individual sets of 25ns trajectories for the data in Figure S14*

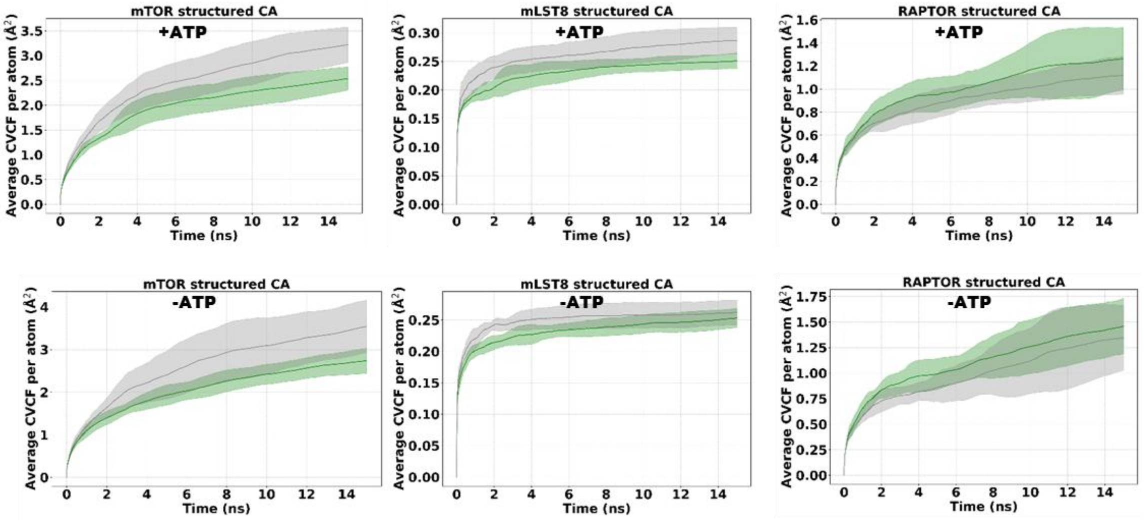


**Figure S16:** *Per atom $\sigma^2_{CVCF}$ traces averages and std-dev over sets of 10×15ns trajectories for the individual protein chains within the four ±RHEB ±ATP mTORC1 complexes.*

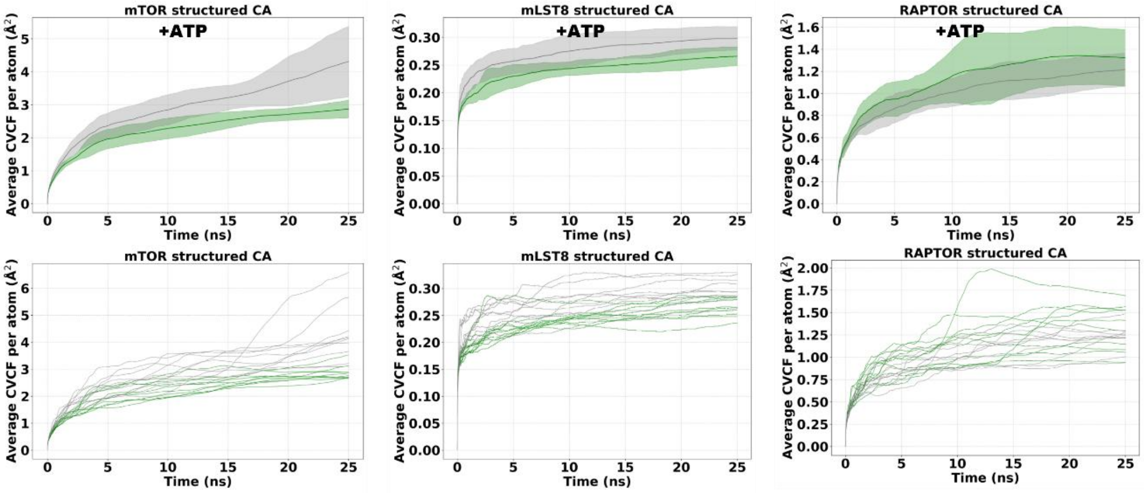


**Figure S17:** (Top row) *Per atom $\sigma^2_{CVCF}$ traces averages and std-dev over sets of 10×25ns trajectories for the individual protein chains within the two ±RHEB mTORC1 complexes with ATP. (Bottom row) $\sigma^2_{CVCF}$ traces for 10 individual trajectories for the data in the top row.*

| PDB ID | Kinase | Atom Pairs | Distance(Å) |
|---|---|---|---|
| 6BCU | mTORC1+RHEB | Asp2338(CG)-ATP(PG) | 7.93 |
| 6BCX | mTORC1-apo | Asp2338(CG)-ATP(PG) | 8.36 |
| 1L3R | cAMPK | Asp166(CG)-ADP-AlF3(Al) | 4.75 |
| 3X2U | cAMPK | Asp166(CG)-ATP(PG) | 6.32 |
| 3X2V | cAMPK | Asp166(CG)-ATP(PG) | 5.64 |
| 3X2W | cAMPK | Asp166(CG)-ATP(PG) | 5.61 |
| 1ATP | cAMPK | Asp166(CG)-ATP(PG) | 5.67 |
| 4DH3 | cAMPKA | Asp166(CG)-ATP(PG) | 5.58 |
| 3FJQ | CAMPK-pep-inh. | Asp166(CG)-ATP(PG) | 5.51 |
| 3QHW | CDK2 | Asp127(CG)-ADP-MgF2(Mg) | 4.8 |
| 1HCK | CDK2 | Asp127(CG)-ATP(PG) | 5.47 |
| 5DRD | Aurora A Kinase | Asp256(CG)-ATP(PG) | 7.2 |
| 4GT3 | ERK2 Kinase | Asp147(CG)-ATP(PG) | 8.34 |
| 1GAG | IRS-Bisubstrate-inh. | Asp1132(CG)-112(PG) | 7.92 |

**Table S6:** *Distance between catalytic Asp with γ-Phosphate/mimic*
